# Building alternative splicing and evolution-aware sequence-structure maps for protein repeats

**DOI:** 10.1101/2023.04.29.538821

**Authors:** Antoine Szatkownik, Diego Javier Zea, Hugues Richard, Elodie Laine

## Abstract

Alternative splicing of repeats in proteins provides a mechanism for rewiring and fine-tuning protein interaction networks. In this work, we developed a robust and versatile method, ASPRING, to identify alternatively spliced protein repeats from gene annotations. ASPRING leverages evolutionary meaningful alternative splicing-aware hierarchical graphs to provide maps between protein repeats sequences and 3D structures. We re-think the definition of repeats by explicitly accounting for transcript diversity across several genes/species. Using a stringent sequence-based similarity criterion, we detected over 5,000 evolutionary conserved repeats by screening virtually all human protein-coding genes and their orthologs across a dozen species. Through a joint analysis of their sequences and structures, we extracted specificity-determining sequence signatures and assessed their implication in experimentally resolved and modelled protein interactions. Our findings demonstrate the widespread alternative usage of protein repeats in modulating protein interactions and open avenues for targeting repeat-mediated interactions.

**Highlights:**

- Robust detection of alternatively used repeated protein regions in evolution
- The approach relies on sequence similarity and identifies conserved signatures
- Mapping of the repeats onto protein isoform 3D models predicted by AlphaFold
- 5 000 repeats detected over the human coding fraction, about one third disordered
- Assessment of the structural coverage of their interactions with protein partners

## Introduction

Repetitive elements are a pervasive feature of proteins, ranging from simple tandem repeats to complex repeat architectures. These repeated regions often play crucial roles in protein function by mediating protein-protein interactions, facilitating protein folding, or acting as flexible linkers between functional domains (Schüler and Bornberg-Bauer, 2016; Kajava, 2012; Andrade et al., 2001). As such, many efforts have been engaged to detect and classify them, as well as characterize their structural and evolutionary properties (Paladin et al., 2021; Delucchi et al., 2020; Pagès and Grudinin, 2019; Schaper et al., 2015; Heger and Holm, 2000; Björklund et al., 2006; Marcotte et al., 1999). Precisely delineating their boundaries remains however challenging. Should it be based on structure, sequence, or a combination of both? A recent census of adjacently repeated sequence patterns, or *tandem repeats*, revealed a wide variability in the characteristics of the repeated units and the number of repetitions (Delucchi et al., 2020). While tandem repeats show enrichment in intrinsic disorder, some fold into specific structures such as solenoids or have “beads on a string” organizations (Kajava, 2012).

Protein repeats are more prevalent in Eukaryotes and Viruses than in Archaea and Bacteria (Delucchi et al., 2020; Marcotte et al., 1999). Toward understanding their origin and expansion, several studies have investigated the extent to which they agree with or violate the exon-intron structure of genes (Paladin et al., 2020; Björklund et al., 2006). These studies uncovered a variety of scenarios, and proposed the use of evolutionary patterns inferred from the intron-exon boundaries across species to refine the delineation and classification of the repeats. Tandem repeats mostly originate through duplication (Delucchi et al., 2020), and exon shuffling can explain their expansion in some families albeit not in all (Björklund et al., 2006).

Recent works have highlighted the importance of alternative splicing (AS) in modulating the number of protein repeats and their amino acid composition (Osmanli et al., 2022; Martinez Gomez et al., 2021; Zea et al., 2021). AS, along with alternative promoter usage and alternative polyadenylation can produce multiple mature mRNA transcripts from a single gene (Ast, 2004). These transcripts may lead to different protein isoforms with distinct shapes (Birzele et al., 2008), interaction partners (Yang et al., 2016), and functions (Baralle and Giudice, 2017; Kelemen et al., 2013). Over the human coding fraction, a couple of thousand genes, mainly involved in cell organization, muscle contraction, and inter-cellular communication, show evidence of evolutionary conserved AS patterns modulating the usage of similar exonic sequences (Zea et al., 2021). Alternative transcripts tend to contain less tandem repeats than canonical ones and the associated deletion events do not alter the overall protein 3D structure (Osmanli et al., 2022). Mutually exclusive tandem homologous exons have drawn the most attention, with substantial efforts for manually curating their annotations and verifying them with transcriptomics and proteomics data (Abascal et al., 2015; Martinez Gomez et al., 2021; Abascal et al., 2015). They are prevalent, have clinical importance, and ancient evolutionary origin.

One key question is defining the pertinent entities to assess the alternative usage of protein repeats. Ideally, these entities should make sense across evolution and account for the full protein diversity generated by AS. Understanding the modularity of protein repeats is also essential, as it may shed light on their functional relevance in protein-protein interactions. We recently proposed a solution for matching protein isoforms across species toward comprehensively describing AS-induced diversity and assessing its evolutionary conservation (Zea et al., 2021). To do so, we introduced evolutionary splicing graphs (ESGs) as a generalization of the notion of splicing graph (Heber et al., 2002) to many genes/species. Each node in an ESG is a spliced-exon, or s-exon, defined as a set of aligned exonic sequences coming from several orthologous genes (**Fig. 1**, top left panel). The s-exons are minimal building blocks for transcripts in evolution.

**Figure 1:**
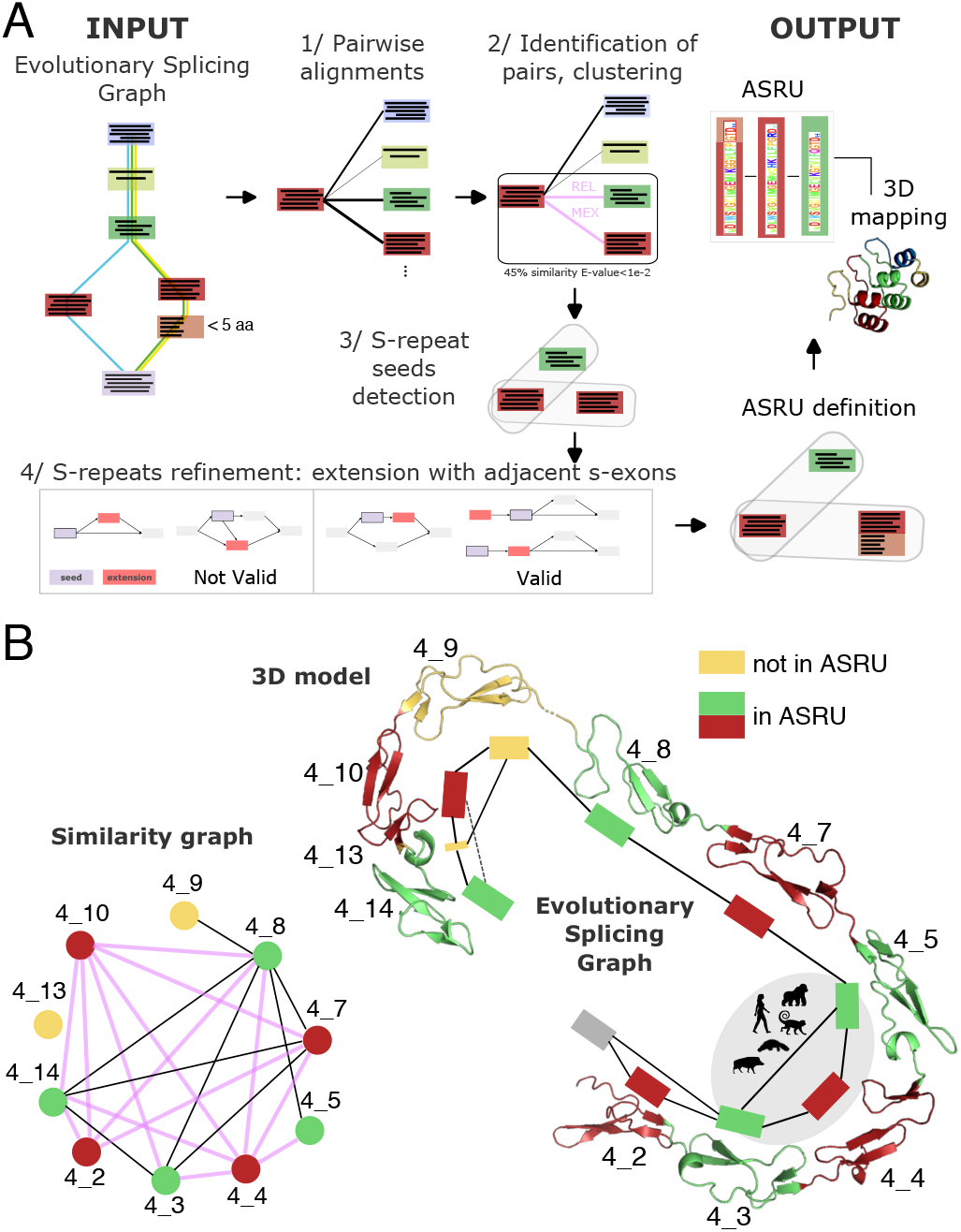
Overview of ASPRING. **A.** Toy example illustrating ASPRING algorithm. In the input ESG, the MSAs defining the s-exons are depicted within the nodes, and three transcripts are highlighted as colored paths. The output ASRU comprises three s-repeats, two of them containing a single s-exon (in red and green) and the third one containing two s-exons (in red and pink). **B**. ASPRING results obtained for Matrilin 2 (**Table S1**). The displayed protein region encompasses one ASRU, made of 8 single-s-exon s-repeats (in red and green) and 2 other s-exons (in yellow). Left panel: S-exon similarity graph, where the s-exons sharing a significantly high sequence similarity are linked by an edge. The pink color indicates an AS relationship. Right panel: 3D model and ESG. The ASRU is associated with three events (solid line bubbles). In the most conserved event, highlighted in grey, both canonical and alternative subpaths are found in primates, platypus and boar. The dotted line indicates another event taking place in this region but not associated with the ASRU, because it does not induce the alternative usage of a repeat. The s-exon *4 9* is not part of the ASRU because, although it shares high similarity with *4 8*, we cannot establish an AS relationship between them (**Fig. S1B**).

In this work, we report on a robust and versatile method, ASPRING (Alternatively Spliced Pseudo Repeats IN-Gene), to identify protein repeats across a set of genes/species with evidence of evolutionary conserved alternative usage. It identifies repeats based on sequence similarity and on the topology of the ESG. By applying ASPRING to the human protein coding fraction, we detected over 5,000 repeats defined across a dozen species spanning 800 million years of evolution. We extracted specificity-determining sequence signatures, providing novel insights into the mechanisms of protein-protein interactions. To further elucidate the functional and structural implications of these repeats, we mapped them onto experimental 3D structures of monomers and complexes available in the Protein Data Bank (PDB) (Berman et al., 2000), as well as 3D models predicted by AlphaFold2 (Burke et al., 2023; Sommer et al., 2022; Varadi et al., 2022; Jumper et al., 2021). Overall, this comprehensive analysis demonstrates the importance of considering alternative splicing patterns in these regions. By identifying specificity-determining sequence signatures and investigating their 3D structure, this study sheds new light on the mechanisms of protein-protein interactions and paves the way for targeted design of repeat-mediated interactions.

ASPRING is freely available at https://github.com/PhyloSofS-Team/aspring and accessible to all as a Python package and a Docker image. Curated data associated with the study are available at https://doi.org/10.6084/m9.figshare.22722682.

## Methods

### Overview of ASPRING

Our approach relies on sequence-based duplication detection and leverages both evolutionary conservation and AS-generated diversity through the use of ESGs (**Fig. 1A**). It first detects pairs of s-exons with similar sequence profiles and then combines them following the topology of the ESG to delineate spliced-repeats, or s-repeats. ASPRING finally groups together similar s-repeats, by transitivity, into alternatively spliced repetitive units (ASRU, **Fig. 1A**, top right). An s-repeat can be viewed as an instance of an ASRU and it is linked to other s-repeats within the same ASRU by sequence similarity and also by a set of AS events. The original contribution of our approach is twofold. It explicitly accounts for both evolution and AS right from the beginning by directly operating on s-exons, instead of species-specific protein sequences. And it defines evolutionary meaningful and AS-aware entities, the ASRUs, thereby going beyond repeat enumeration.

ASPRING takes as input a Gene Ensembl ID and an evolutionary splicing graph (ESG) computed by ThorAxe (Zea et al., 2021) for this gene and its one-to-one orthologs in a set of species. The ESG summarizes the transcript diversity observed in this set of orthologous genes (**Fig. 1A**, top left panel). Each node in the ESG, called a sexon, is an alignment of translated exonic sequences coming from different genes/species. An edge between two nodes indicates that the corresponding s-exons co-occur and are consecutive in at least one observed transcript isoform. Hence, the s-exons can be viewed as the minimal building blocks of the isoforms in evolution. Starting from the input ESG, ASPRING extracts a set of ASRUs (**Fig. 1A**, top right panel). Each ASRU is defined by a collection of s-repeats, where an s-repeat can be a single s-exon or an ordered list of several s-exons showing sequence similarity and evidence of alternative usage. ASPRING outputs several tables summarizing information about the ASRUs, the s-repeats, the similar pairs of s-exons and the raw alignments, and also annotated 3D models of the protein isoforms corresponding to the input transcripts (**Fig. 1A**, top right panel).

ASPRING algorithm unfolds into four main steps (**Fig. 1A**, see also *Supplementary Methods*). It first performs an all-to-all comparison of the s-exons in the input ESG using profile HMMs with the HH-suite3 (Steinegger et al., 2019) (**Fig. 1A**, step 1). Secondly, it identifies similar s-exon pairs by considering four criteria, namely the significance of the p-value associated with the profile HMM-HMM alignment, the sequence identity and the coverage shared between the s-exons, and their evolutionary conservation (**Fig. 1A**, step 2). By default, we require a high level of shared sequence identity (45%) to allow for contrasting amino acid divergence across isoforms versus species. All filtering criteria can be changed by the users depending on their system of interest. As this point, since our motivation is to assess the AS-induced modulation of the composition and number of protein repeats, we need to verify the existence of some alternative s-exon usage. For this, ASPRING explicitly accounts for the AS events encoded in the topology of the ESG.

Following (Zea et al., 2021), we define as AS event as a variation from a reference *canonical* transcript chosen for its high conservation and length (**Fig. S1A**). Given a pair of s-exons, ASPRING considers four types of AS-mediated relationships, namely MEX, ALT, REL and UNREL, inferred from the role of each s-exon in the AS events (**Fig. 1A**, see labels on the pink edges, and **Fig. S1B**). We introduced these four relationships in (Zea et al., 2021). Briefly, MEX s-exons are mutually exclusive and ALT s-exons are interchangeable but not in an exclusive fashion. These two relationships correspond to AS events of the types mutually exclusive and alternative (**Fig. S1**). In a REL pair, one s-exon is located inside the bubble representing an AS event and is thus alternatively used, while the other serves as an anchor for the event and is thus always expressed. The fourth relationship, UNREL, is the weakest one: one s-exon is part of an AS event while the other is located outside the event. The REL and UNREL pairs may be inferred from different types of AS events (**Fig. S1**). In terms of protein sequence proximity, the MEX s-exons are exactly at the same location (and consecutive in the genome). The ALT s-exons can be at the same place or next to each other. The REL s-exons are typically also close to each other in the protein, whereas the UNREL s-exons may be located in remote and completely different contexts. ASPRING determines for each pair of similar s-exons whether it is of type MEX, ALT, REL or UNREL, in that priority order in case multiple relationships exist. It filters out the pairs that do not fit any category, and thus that do not have any evidence of alternative usage.

Next, ASPRING algorithm’s third step clusters the selected s-exons using the transitivity principle (**Fig. 1A**, step 3). For any two valid pairs of s-exons sharing an s-exon in common, *e*.*g. (A,B)* and *(A,C)*, ASPRING will group the three s-exons *A, B* and *C* in the same cluster. The procedure amounts to detecting connected components in a graph where the nodes are the s-exons and the edges indicate valid pairs of similar s-exons. The resulting clusters can be viewed as prototypes of the ASRUs and the s-exons composing them are s-repeat seeds. In the final fourth step, the ASPRING algorithm aims at extending these seeds with adjacent s-exons, according to the topology of the ESG, to obtain more homogeneous ASRUs in terms of length (in aa) of their s-repeats (**Fig. 1A**, step 4). We ensure that the s-exons used for extending the seeds comply with the same sequence similarity constraints as the seeds themselves.

On top of the ASRU detection, ASPRING maps the s-exons and s-repeats onto the 3D models of the input isoforms available from the AlphaFold database (Varadi et al., 2022) (**Fig. 1A**, top right panel). This functionality allows users to easily visualise the sequential and spatial localisation of the s-exons and s-repeats.

### Application over the human coding fraction across a dozen species

We ran ThorAxe (Zea et al., 2021) on the ensemble of 18 228 human protein-coding genes and their one-to-one orthologs across 12 species, namely three primates (*Homo sapiens, Gorilla gorilla, Macaca mulatta*), two rodents (*Rattus norvegicus, Mus musculus*), four other mammals (*Bos taurus, Sus scrofa, Ornithorhynchus anatinus, Monodelphis domestica*), one amphibian (*Xenopus tropicalis*), one fish (*Danio rerio*), and one nematode (*Caenorhabditis elegans*). We downloaded the corresponding gene annotations from Ensembl release 105 (December 2021) (Cunningham et al., 2022). We used the default parameters except that we did not apply any filtering of the transcripts based on Transcript Support Level. On average, each human gene had one-to-one orthologs in 8 of the other species (**Table S2**). About 7% of the genes were found in all 12 species. The detection of ASRUs by ASPRING is conditioned by the presence of AS events in the ESG computed by ThorAxe. The latter detected AS events (supported by at least two transcripts) in a subset of 13 454 genes. We thus ran ASPRING on this subset, using the default parameters (see *Supplementary Methods*). The calculation took approximately 4-5 days on 100 CPUs Intel Xeon Silver 4210 2.2GH. The most computationally expensive step was the generation of the profile HMM alignments with the HH-suite 3 (Steinegger et al., 2019). Once these alignments have been computed for a given gene, the users can re-run ASPRING with different filtering criteria within a few minutes.

We performed a comprehensive analysis of the evolutionary signals encoded in the detected repeats, their 3D structural properties and interactions. For the evolutionary analysis, we focused on the information contained in the alignments generated by the HH-suite. For the structural analysis, we predicted disorder using the protein Language Model-based method SETH (Ilzhoefer et al., 2022). To characterize interactions, we looked at all the physiologically relevant experimental 3D complex structures involving the ASRU-containing proteins or their close homologs. We detected interfaces based on a distance criterion and we mapped them to the sequences of the s-repeats. Experimental PDB structures are often partial and may contain modifications, which makes the mapping sometimes difficult. To circumvent this issue, we used the full-length transcript 3D models available from the isoform.io database (Sommer et al., 2022). In addition, we explored a resource of 3D complex models predicted by AlphaFold2 Burke et al. (2023). See Supplementary methods for detailed protocols.

## Results and Discussion

### ASPRING provides AS- and evolution-aware sequence-structure maps for repeats at the human proteome scale

We used ASPRING to uncover the alternative usage of protein repeats on the human proteome scale. We analysed 18 228 human protein-coding genes and their orthologs across 12 species spanning 800 million years of evolution, from primates to nematode. In total, ASPRING screened over 8 million s-exon pairs, among which about 26 500 passed all four filters for p-value, identity, coverage and conservation (**Fig. S2**). We applied a stringent sequence identity threshold of 45% to allow for contrasting the divergence between repeats coming from the same species versus the cross-species divergence of orthologous repeats. Among the retained s-exon pairs, about 11 000 are linked by some alternative splicing event. ASPRING grouped these s-exon pairs in 1 469 alternatively spliced repetitive units (ASRUs) coming from 1 039 genes. About 7% of these genes have one-to-one orthologs in all 12 species, in line with the proportion computed over the whole proteome (see *Methods*), and they contribute 73 ASRUs. An ASRU is made of at least two s-repeats, each one corresponding to a single s-exon or several consecutive s-exons (see *Methods*). In general, the longer the gene the higher the number of detected ASRUs (**Fig. S3**). The s-repeats span a broad range of lengths, from 5 to over 5 000 amino acid residues, with a median of 36 residues. Despite this variability, most of the ASRUs display a strong regularity in their s-repeat lengths (**Fig. S4**). We often observed that some s-repeats are exact multiples of others, with multiple similarity hits (**Fig. S4**).

Matrilin 2 shows an illustrative example of the AS- and evolution-aware maps produced by ASPRING (**Fig. 1B**). In this protein, we found one ASRU defined by an array of 8 s-repeats (in red and green) of very regular lengths, namely 41±5 residues. Within this array, three s-repeats are alternatively included or excluded, namely *4_2, 4_4* and *4_10*. The most conserved event involves *4_4* and is found in primates, platypus and boar.

Some other s-exons are located within the array but do not belong to the detected ASRU. For instance, while the s-exon *4_9* could be considered as a repeat based on its similarity with *4_8*, we cannot establish any AS relationship between these two s-exons (see **Figure S1B**). Likewise, the event defining the alternative inclusion/exclusion of the s-exon *4_13*, although located in the same protein region, is not associated with the ASRU because it does not induce the alternative usage of any repeat.

Overall, about two thirds of the 1 039 ASRU-containing genes have exactly one ASRU made of two s-repeats (**Fig. 2A**), in line with previous observations (Osmanli et al., 2022). Examples include the actinin *α* chains, the integrin subunit *β*1, the Ras-related proteins Rab-37 and Rab-6A, several Solute Carrier Transporters, the MAP kinases 8, 9, 10 and 14, and the synaptosome associated protein 25, where ASPRING automatically detected pairs of alternatively spliced tandem duplicated exons previously reported through manual curation (Martinez Gomez et al., 2021). These pairs have proteomic evidence, clinical importance and ancient evolutionary origin (Martinez Gomez et al., 2021). In a couple of hundred genes, the detected ASRUs contain many s-repeats (**Fig. 2A**), up to 127 for the giant skeletal muscle protein Nebulin for instance. They often correspond to known tandem repeats detected in a row (**Fig. 2B**, on top, in green). ASPRING is however not limited to detecting tandem repeats and some ASRUs contain repeats lying at distant locations in the protein (**Fig. 2B**, on top, in orange). Among the proteins with the most populated ASRUs, we found giant chains playing structural roles, such as those from the collagen. Finally, a minority of genes contain many ASRUs (up to 35), each one made of a few s-repeats (**Fig. 2A**). In such cases, the sequence similarity is at the level of entire domains and the different ASRUs correspond to different parts of these domains (**Fig. 2B**, at the bottom). ATPases, platelet glycoprotein 4, sucrase-isomaltase, and ABC transporters partake in this archetypal situation.

**Figure 2:**
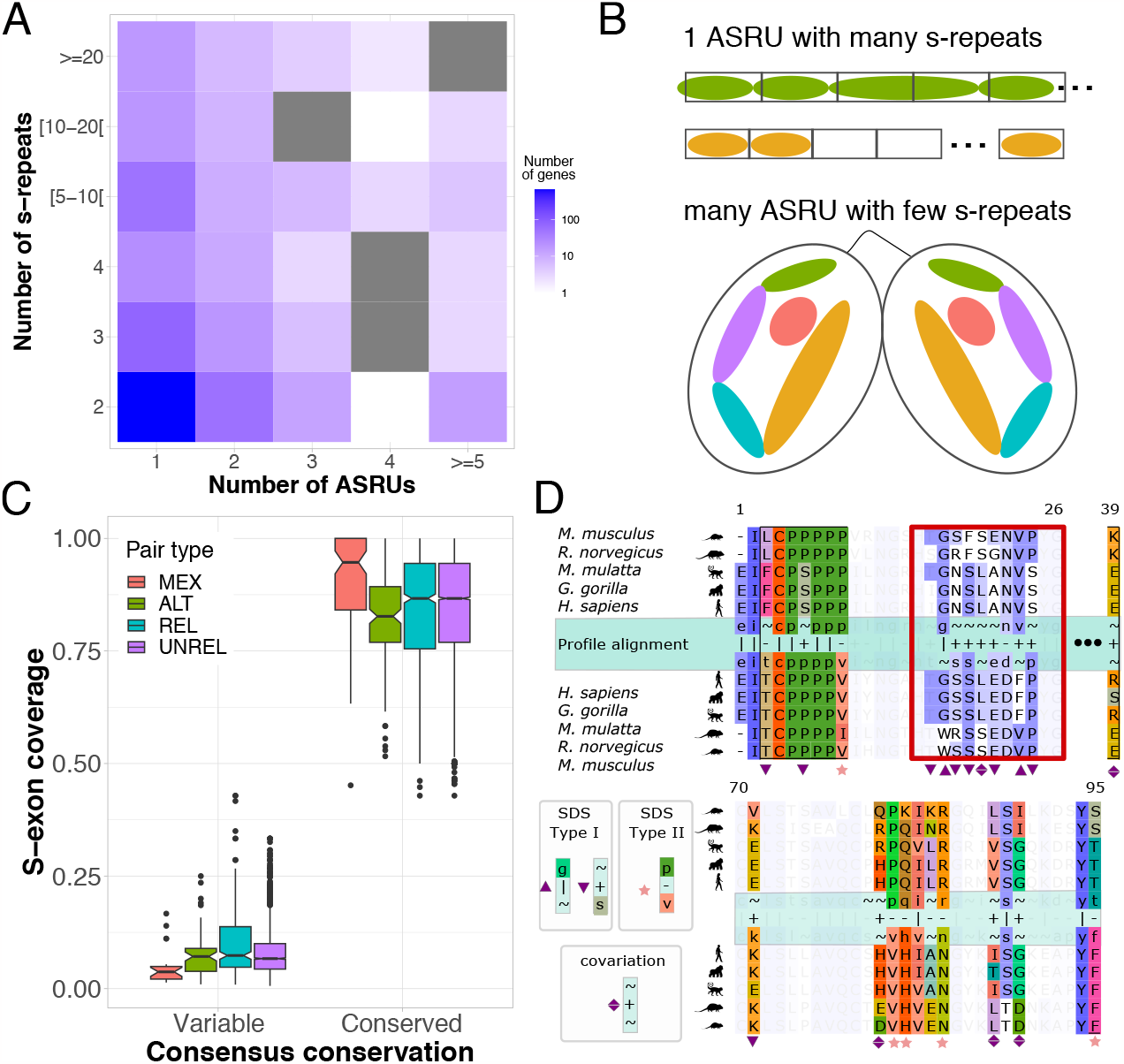
Overview of the landscape of alternatively used repeats in the human proteome. **A.** Heatmap showing the distribution of ASRUs and their s-repeat composition. Each cell (*x, y*) gives the number of genes with *x* ASRUs and whose largest ASRU contains *y* s-repeats. The color scale is in logarithm and grey cells are not populated. **B**. Three Archetypal situations. On top, 1 ASRU made of many tandem repeats detected in a row (in green) or repeats located at more distant locations in the protein (in orange). The s-repeat boundaries approximately match those of the known repeats (black rectangles). At the bottom, many ASRUs (each colored differently) made of a couple of s-repeats coming from two domains (delineated in black). **C**. Distributions of the proportion of aligned positions between two s-exons that are variable or conserved. Variable (respectively, conserved) positions have their major amino acids conserved in less (resp. more) than 40% of the species within each s-exon. **D**. Detection of sequence signatures (gene CR2). The alignment between the profile HMM for s-exons 1 2 and 1 3 is shown in the middle (green band), together with the MSA for each of the s-exons. For each pair of s-repeats, we analyse the profile-profile alignment to detect positions that harbour different alignment signatures. Each possible signature is based on the degree of conservation within the s-repeat (∽: not conserved), and on the score of the aligned position between the s-repeats. Three types of signatures are detailed in the figure: SDS Type I, SDS Type II, and covariation.

### ASPRING unveils evolutionary ancient patterns of repeat modulation

The vast majority of the s-exons in the detected s-repeats are highly conserved, as measured by the *species fraction* that is the proportion of species where a s-exon is found (**Fig. S5A**). Regarding the AS events, mutually exclusive and alternative events are enriched within ASRUs compared to the proportions computed over the entire proteome (**Fig. S5A**). By contrast, insertions are under-represented, in agreement with a recent study (Osmanli et al., 2022). This observation indicates that the s-repeats are often considered as part of the canonical transcript by ThorAxe and their alternative inclusion or exclusion appear as deletions with respect to that transcript. It underlines the high evolutionary conservation of the s-repeats, since the canonical transcript is chosen as the most represented across species (see *Supplementary Methods*). Moreover, the s-repeat junctions in the ESG tend to be well preserved between human and the other species (**Fig. S5C**). On average, the s-repeats conserved across Eutheria have more than 75% of their junctions preserved (**Fig. S5C**). This proportion is still higher than 70% in Metatheria (opossum), Prototheria (platypus) and Amphibia (frog), and higher than 65% in Teleostei (zebrafish). Overall, the s-repeat junctions are better preserved than the global topology of the ESG (**Fig. S5D**).

An AS event is defined by a couple of subpaths in the ESG, one being canonical and the other alternative. We found that within each species, most of the events linking the s-repeats are supported by only one of the two subpaths (**Fig. S6A**). For instance, only 30% of the 489 events with both the canonical and alternative subpaths annotated in human also have both subpaths annotated in mouse (**Fig. S6B**). This result reveals that the alternative usage of the repeats is spread across the different species. Nevertheless, for a substantial amount of events, both subpaths are found in several species, revealing evolutionary ancient modulation patterns in the usage of protein repeats. For instance, 89 events are well represented across mammals (at least 6 out of 9 species). A subset of 64 events are conserved from primates to amphibians, among which 14 extend to teleosts (**Fig. S6B**). These events concern ASRUs detected in the proteins CACNA1C, CAMK2D, CDC42, EYA4, FGFR2, IDH3B, KRAS, OTOF, PITPNB, RAB6A, RELCH, and SLC39A14, all previously reported as functionally and clinically important (Martinez Gomez et al., 2021), and two newly identified ASRUs in JAKMIP1 and TNIK. Finally, each s-repeat within an ASRU may or may not be directly impacted by the events. For the ASRUs made of only two s-repeats, in the most frequent scenario, namely 2/3 of the cases, both s-repeats are subject to alternative inclusion/exclusion. For the bigger ASRUs, we observed a variety of scenarios, from a few specific s-repeats being alternatively spliced to the full s-repeat array (**Fig. S6C**). For example, in Matrilin 2, three out of eight s-repeats are modulated (**Fig. 1B**), whereas in Nebulin, 90% of the 127 s-repeats forming the biggest ASRU are subject to alternative inclusion or exclusion through 47 events spread across species. Overall, the alternatively spliced s-repeats tend to be less conserved than the constitutive ones (**Fig. S6D**).

### Evolutionary signals suggest functional diversification of the repeats

We asked whether the s-repeats belonging to the same ASRU are subject to the same evolutionary constraints. To do so, we adopted a human-centred perspective by focusing on the s-exons and s-repeats present in humans and by taking the human exonic sequences as references. We systematically assessed and compared the conservation profiles of the s-exons ASPRING used to build the ASRUs (see *Materials and Methods*). We found that the vast majority of the positions in the s-exon pair alignments are highly conserved (**Fig. 2C**). More precisely, both s-exons in the pair feature a conserved consensus amino acid (aa) at 85% of the aligned positions, on average. The mutually exclusive pairs display the strongest signal (**Fig. 2C**, MEX), with 90% of their aligned positions conserved in both s-exons. This observation agrees with previous studies emphasising the ancient evolutionary origin and functional importance of such pairs (Martinez Gomez et al., 2021; Abascal et al., 2015). Less than 10% of the positions are variable in both s-exons (**Fig. 2C**).

The s-repeats belonging to the same ASRU are mostly indistinguishable, often because they share the same highly conserved aas (79% of the conserved aligned positions) and in a few cases because they display highly variable and unrelated sets of aas (87% of the remaining positions). Nevertheless, some clear-cut patterns suggest functional diversification of the s-repeats. We illustrate these patterns with the two s-repeats detected in the complement C3d receptor 2 (**Fig. 2D**). At positions marked with a pink star, the same strong selection pressure applies to both s-repeats but for different aas. They correspond to type II specificity-determining site (SDS) in (Chakraborty and Chakrabarti, 2015). For instance, at position 82, the two s-repeats have highly conserved aas exhibiting distinct physicochemical properties, namely Proline and Valine. The same type of pattern occurs at position 9, with the minor difference that Isoleucine replaces Valine in one species, these aas being very similar. This type II SDS contrasts with the preceding positions 2, 4, 5, 7, and 8, where the s-repeats share the same highly conserved aas. Positions 83, 86, and 98 also qualify as type II SDSs, albeit with less striking physicochemical differences or weaker conservation. Positions highlighted with purple triangles indicate a redistribution of the selection pressure over the s-repeat sequences (**Fig. 2D**). They correspond to type I SDS, conserved in one s-repeat and variable in another (Chakraborty and Chakrabarti, 2015).

Beyond these strong conservation signals, several variable positions display non-random covariation patterns suggesting that the same selection pressure has applied to the s-repeats despite phylogenetic divergence (**Fig. 2D**, purple diamonds). For instance, at positions 20 and 87, the two s-repeats are always in match, although the sequences have diverged between rodents and primates. Namely, the Leucine observed in primates at position 20 corresponds to a Serine in rodents. Position 39 features a persistent charge pattern, despite sequence divergence. More precisely, in primates, the first s-repeat displays the negatively charged Glutamate while the second one has the positively charged Arginine. Reversely in rodents, the second s-repeat carries the negative charge and the first s-repeat has the positive one (Lysine).

Overall, we identified 856 ASRUs coming from 625 genes with SDS or covariation sites. They amount to almost two thirds of all ASRUs detected by ASPRING, and most of the time, they contain more than just one type of sites (**Fig. S7A**). The s-exon pairs that do not contain any SDS or covariation sites tend to have more highly conserved positions with a shared amino acid (perfect match) and more weak matches (**Fig. S7B**).

### 3D structural properties and interactions of the repeats

About two thirds of the s-repeats were predicted to fold into a stable 3D structure by SETH, based on their sequences (see *Methods*), and about one third was predicted as disordered (**Fig. 3A**). The prediction is generally consistent over the full length of the s-repeat. In other words, either a s-repeat is fully ordered, or it is fully disordered, with very few in-between cases (**Fig. 3A**). This highly bimodal distribution indicates that the s-repeat boundaries tend to agree with structural elements boundaries. Regarding sequence divergence, we did not observe any dependence of the quality of the MSAs on the tendency to fold (**Fig. 3A**, inset).

**Figure 3:**
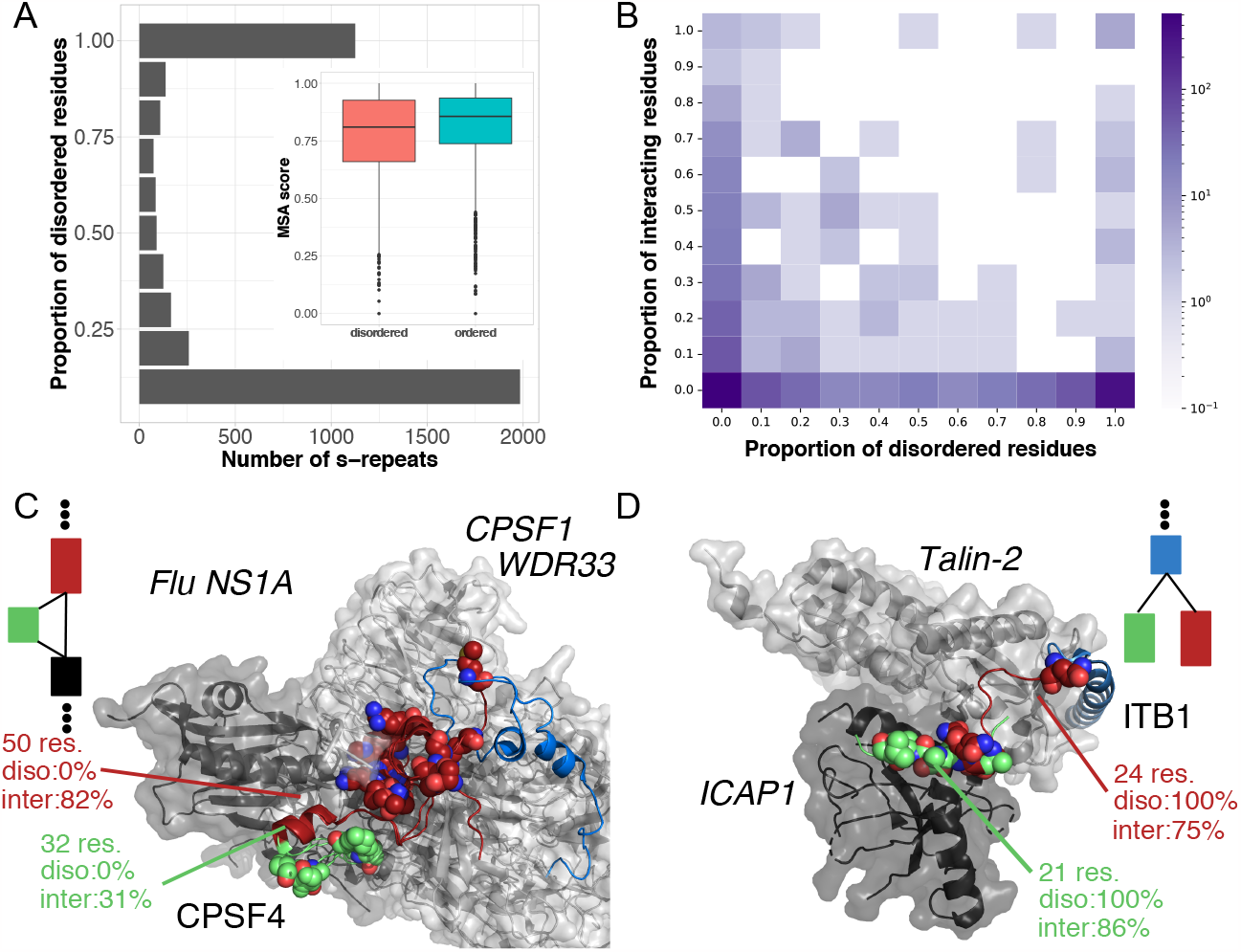
3D structures and interactions of the s-repeats. **A.** Distribution of the content of intrinsic disorder predicted in the s-repeats (see *Supplementary methods*). The plot in the inset compares the distributions of MSA scores for the disordered (salmon) versus ordered (turquoise) s-repeats. **B**. Heatmap of the proportion of s-repeat residues involved in the physical interaction with a partner versus the proportion of residues predicted as disordered. We considered only the s-repeats belonging to a protein with experimental structural information (see *Supplementary methods*). **C-D**. Examples of s-repeats in interaction with protein partners. In both cases, the ASRU-containing protein is displayed as cartoons and the protein partners are displayed as transparent surfaces, with their names highlighted in italics. The s-repeats are highlighted in firebrick and lime. Type II SDS are indicated by spheres. The proportion of residues predicted as disordered and of residues in interaction are given for the two s-repeats, along with their lengths. A portion of the ESG showing the AS event linking the two s-repeats is also depicted. **C**. PDB codes 2RHK and 6URO for the complexes of CPSF4 bound to Flu NS1A and CPSF1-WDR33, respectively. **D**. PDB codes 3G9W and 4DX9 for the complexes of ITB1 bound to Talin-2 and ICAP1, respectively.

For a third of the s-repeats, we could cross the disorder prediction with experimental structural information from the PDB about their interactions with protein partners (**Fig. 3B**). In total, we found 351 s-repeats in direct physical contact with another protein in a physiologically relevant macromolecular 3D complex. As expected, most of these s-repeats, namely 75%, were predicted as fully ordered by SETH, *i*.*e*. they have less than 20% of disordered residues (**Fig. 3B**). The cleavage and polyadenylation specificity factor subunit 4 (CPSF4) gives an example of two consecutive s-repeats predicted as fully ordered and participating in the interfaces with two partners, namely the influenza NS1A protein and the pre-mRNA 3’-end processing complex (**Fig. 3C**). These interfaces comprise twelve type II SDS. The two s-repeats have ancient evolutionary origin, *i*.*e* they are highly conserved from human to zebrafish. They encompass the four first Zinc-finger domains, Zn-1, Zn-2, Zn-3 and Zn-4 annotated in CPSF4. While the larger s-repeat is always expressed, the smaller one is deleted in some transcripts (REL relationship). This deletion, conserved in human, gorilla, mouse, opossum, platypus and frog, results in a shift of the Zinc fingers, with Zn-4(Cter)-Zn-5(Nter) taking the place of Zn-3(Cter)-Zn-4(Nter), thus substantially modifying the composition of the interacting surface (**Fig. 3C**). This observation suggests a direct impact of the AS event on the binding affinity.

Ordered s-repeats are not the only ones to mediate interactions, and we found that about 10% of the 351 interacting s-repeats we identified were predicted as fully disordered (*>*80% of their residues, **Fig. 3B**). This result suggests a significant contribution of disorder-to-order transition upon protein repeat binding to their partners. In particular, the integrin *β*1 (ITB1) features an ASRU made of two mutually exclusive tandem s-repeats predicted as fully disordered, and yet, resolved in complex with two partners, namely Talin-2 and ICAP1, respectively (**Fig. 3D**). ITB1 is one of the few proteins for which the 3D structures of several splice variants have been experimentally characterised. Even more, the impact of alternative splicing on the binding affinity between ITB1 and Talin-2 has been assessed (Anthis et al., 2009). More specifically the isoform displaying the s-repeat *18_0* has higher affinity for Talin-2 than that displaying *18 1*, and this higher affinity has functional consequences in myotendinous junctions (Anthis et al., 2009). Both s-repeats have ancient evolutionary origin and their alternative usage is conserved from primates (human, gorilla and macaque) through mammals (rat, boar, cow, opossum, platypus) to amphibians (frog). Several residues differing between the two s-repeats and fully conserved across species are part of the interacting surface, suggesting a direct implication in modulating binding affinity. The microtubule-associated protein tau (MAPT), involved in neurodegenerative diseases, gives another example of disordered s-repeats resolved in 3D complex structures. The ASRU detected in this protein is made of four s-repeats *9_0, 8_0, 8_1, 17_0* that matches well the array of four tandem repeats described in the literature and forming the MT-binding domain. Although intrinsically disordered, tau can adopt a “paperclip” conformation where the MT-binding domain and the N- and C-terminus interact (Strang et al., 2019; Jeganathan et al., 2006).

Out of the *∼*90 000 possible binary interactions between the ASRU-containing proteins that have a 3D structure in the PDB, 248 are supported by a physiologically relevant experimental complex structure (**Fig. S8**). A couple of proteins, namely Polyubiquitin-C and Calmodulin, have more than 3 partners in the set. To extend the scope of the analysis, we considered the 60 000 AlphaFold-predicted binary complex models (Burke et al., 2023) available for the Human Reference Interactome (HuRI) (Luck et al., 2020). These interactions have been experimentally identified in Human. The ASRU-containing proteins detected by ASPRING are involved in a bit less than 10% of these complexes. Each 3D model is associated with a predicted DockQ score, a continuous measure of protein-protein complex model quality (Basu and Wallner, 2016). We retained only the models with a predicted DockQ score (pDockQ) higher than 0.23. At this threshold, 70% of the complexes are expected to be correctly modelled (Burke et al., 2023). We identified 55 models representing an interaction between two ASRU-containing proteins (heterodimer) or between two copies of the same ASRU-containing protein (homodimer, **Fig. 4**). In essentially all these complexes, some s-repeats are structured and take part in the binding interface. Nevertheless, in most of the heterodimers, *e*.*g*. between keratin 78 and keratin 16, the structured interfacial s-repeats come from only one partner, the s-repeats of the other partner being modelled with low confidence.

**Figure 4:**
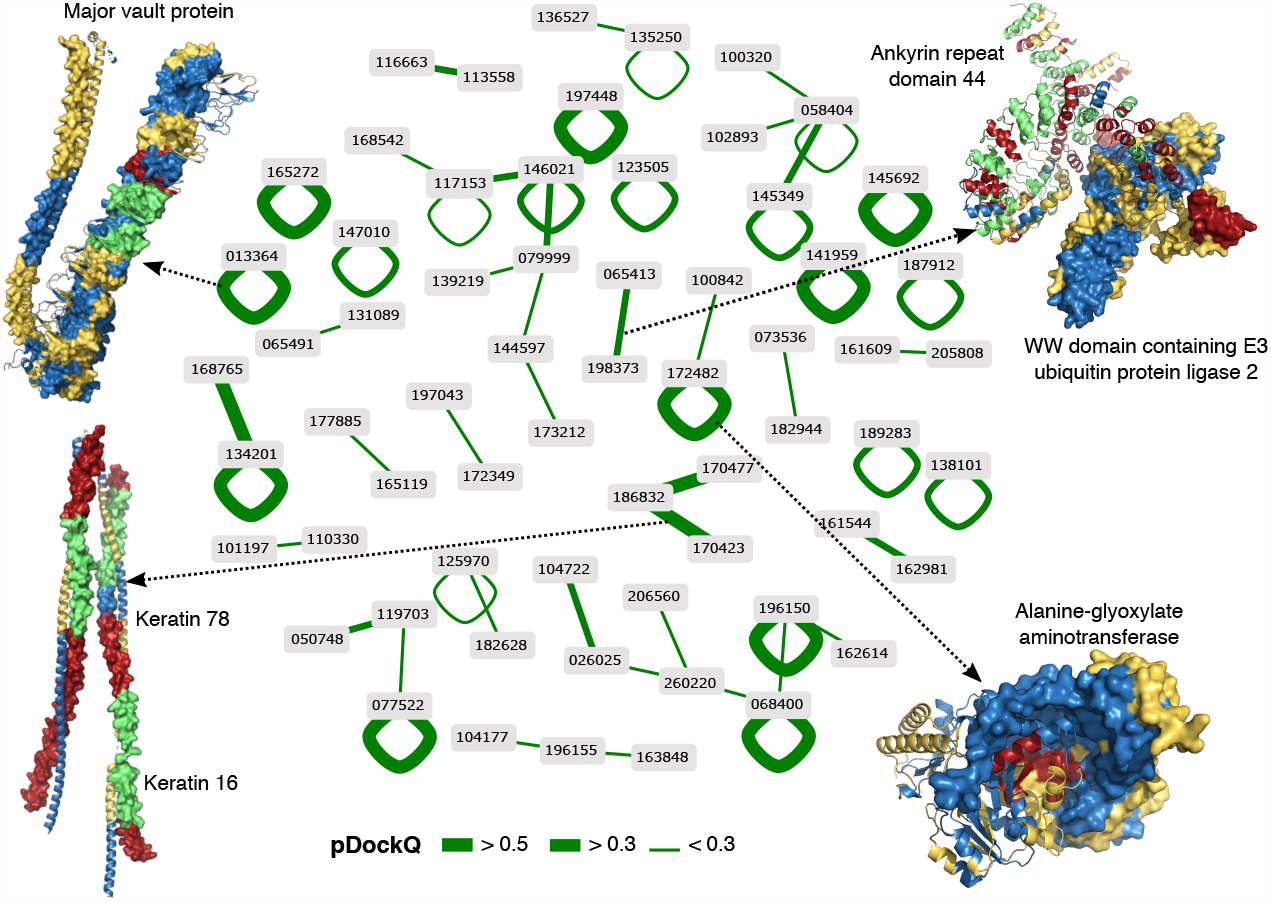
AlphaFold-predicted 3D interaction network among the ASRU-containing proteins. Each node represents an ASRU-containing gene and is labelled with its Ensembl id (prefix: ENSG00000). Each edge represents an interaction experimentally verified in (Luck et al., 2020) and structurally modelled in (Burke et al., 2023). For each protein, Burke and co-authors considered only the main isoform annotated in Uniprot (Burke et al., 2023). The thickness of the edges indicates the predicted model quality (pDockQ score). On the illustrative 3D structures, one partner is shown as cartoons and the other one as surface. The low-confidence regions (pLDDT*<*0.5) are not shown. The s-repeats detected by ASPRING are alternatively colored in green and fire-brick while the remaining s-exons are in blue and yellow. The network was rendered with LEVELNET (Behbahani et al., 2023).

### Comparison with known structure-based tandem repeats

We tested whether ASPRING could detect structure-based repeats annotated in RepeatsDB (Paladin et al., 2021). We selected a set of 10 proteins for which the mapping between structural repeats and exons has been extensively described in (Paladin et al., 2020). By lowering down the sequence identity cutoff at 30%, ASPRING could detect ASRUs in three ankyrin repeat (ANK) and leucine rich-repeat (LRR) containing proteins (**Fig. 5**). The ASRU detected in human ankyrin-1 comprises 19 s-repeats, all highly conserved from human to frog (**Fig. 5A**). Ten of the s-repeats have exactly the length of a classical Ankyrin unit (33 residues, highlighted by one triangle). Two of them have double length (66 residues, highlighted by two triangles) and span two units. These observations are in line with those reported in (Paladin et al., 2020). Deviations from these classical patterns are visible at the extremities, and also in the middle of the protein, where two s-repeats, each made of two s-exons, namely *9_13-9_14* and *9_16-9_17*, are alternatively included or excluded in some transcripts from rat and frog (**Fig. 5A**, ESG on top). The relatively small (16 residue-long) s-exon *9_15* (in blue) serves as an anchor for these two AS events. In murine tankyrase-1, the detected ASRU contains 16 s-repeats, all highly conserved from human to frog and comprising each a single s-exon (**Fig. 5B**). In the experimentally resolved part of the protein (rectangle) we observed that the s-exon boundaries match well the structure-based repeats annotated in RepeatsDB, in agreement with (Paladin et al., 2020). The AlphaFold model allows substantially expanding the structural coverage of the protein, uncovering previously unresolved repeats. The AS event linking the s-repeats substitutes the 744 C-terminal residues of the protein, encompassing 6 s-repeats, by a 34-long unrelated segment. The human leucine rich-repeat (LRR) containing G-protein coupled receptor 5 features ten single-s-exon s-repeats, conserved from human to frog (**Fig. 5C**). They encompass up to 3 structural repeats annotated in RepeatsDB, most of them (8/10) matching well a single structural repeat. As noted in (Paladin et al., 2020), we observe a very regular pattern, with s-repeat boundaries slightly shifted (by 2-3 amino acids) compared to the structural repeats. Some of the later (5 out of 18 are not included in the ASRU due to their high sequence divergence. The number of s-repeats is modulated by three deletion events, two of them conserved across species and one observed only in frog. In all three proteins, the s-repeats display several type I and type II SDS. In the other solenoids and *β*-propellers emphasized in (Paladin et al., 2020), ASPRING did not detect any ASRU due to a low sequence similarity between the structural repeats or a lack of evidence for alternative usage. For instance, in the *β*-propellers, only a fraction of the repeats share more than 30% sequence similarity, challenging their exhaustive detection. This issue may be circumvented by lowering down the sequence identity threshold. Nevertheless, if these repeats are not alternatively used across species, then ASPRING will not consider them further. For example, in the LRR-containing protein RNH1, ASPRING detected 19 similar s-exon pairs encompassing the annotated structural repeats but they were not linked by any AS event. We observed a similar scenario for the two other solenoids Pumilio-rich protein PUM1 and the HEAT-containing protein XPO1.

**Figure 5:**
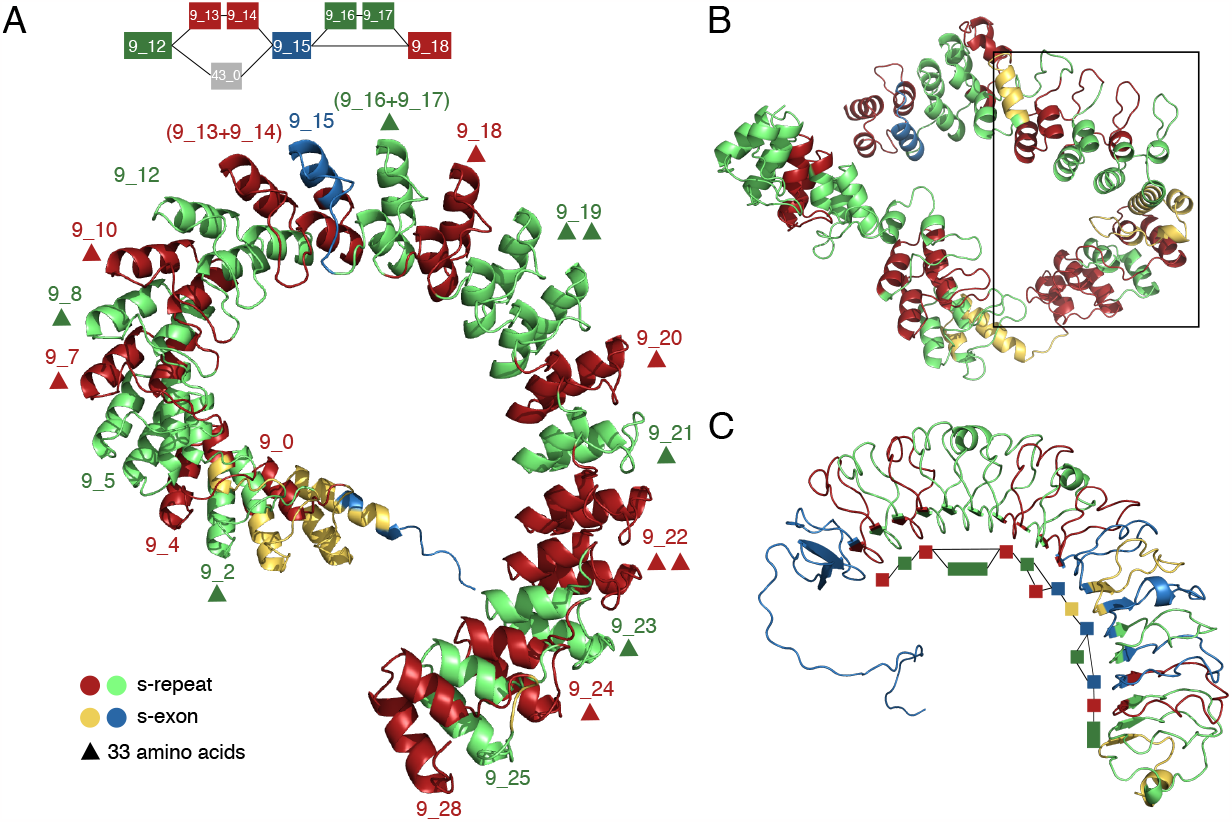
Detection of alternatively spliced repetitive units in solenoids. The s-repeats detected by ASPRING are mapped onto AlphaFold 3D models and are highlighted in green and red. The s-exons that are not part of a s-repeat are colored in blue and yellow. For ease of visualisation, we used an alternating coloring scheme. The analysis was performed on groups of orthologous genes across 12 species, from human to nematode. **A**. Human ankyrin 1. A portion of the ESG is shown on top, with the coloring matching that of the 3D model. **B**. Murine tankyrase-1. The structural repeats annotated in RepeatsDB are highlighted with a black rectangle. **C**. Human leucine rich-repeat (LRR) containing G-protein coupled receptor 5 (LGR5). The ESG is displayed with a layout following the 3D structure, and using the same coloring. Identifiers and residue spans are given in **Table S1**.

## Conclusion

This work proposes an automated method for detecting alternatively used protein repeats through the prism of alternative splicing in evolution. In contrast to other works, we do not aim at exhaustively enumerating repeats but rather at guiding the users to focus on protein “units” sharing complex relationships, namely sequence similarity, conservation in evolution, and alternative usage. Our method is highly versatile, allowing the users to adapt their experimental setup depending on the level of similarity and of conservation they expect or target. For instance, in the case of structural repeats, it may be beneficial to lower down the sequence identity and p-value cutoffs. This is an easy and inexpensive task that users can perform iteratively when analyzing a particular protein since the pipeline does not need to realign the s-exons for testing them. Nevertheless, the current approach might not be adapted for cases where sequence similarity is extremely low. A future improvement could be to allow for performing the similar s-exon pair detection step relying on structure similarity, instead of sequence similarity, for instance using Foldseek (van Kempen et al., 2023), or other methods. We applied ASPRING at large scale over hundreds of million years of evolution. We found that the detected repeats tend to be evolutionary conserved and we identified evolutionary ancient modulation patterns in their usage. We relied solely on gene annotations from the Ensembl database. We expect such annotations to grow rapidly in the coming years, thanks to the advent of long-read sequencing technologies. In this context, methods such as ASPRING will become instrumental to shed light on protein diversification. Future work will focus on expanding the detection to paralogous families and integrating raw RNA-Seq data. Our joint analysis of the sequence patterns, structural properties and interaction propensities exhibited by the repeats, at the amino acid resolution, provides a first step toward improving our understanding of how AS-induced variations modulate the shape, the strength, the stoichiometry and the specificity of repeat-mediated protein interactions. We identified and quantified specificity determining sites in the detected repeats. We estimated the extent of intrinsic disorder and experimentally resolved or modelled interaction surfaces. We showcased the putative role of sequence signatures in establishing and stabilising physical interactions in a few examples. We showed that our approach allows automatically detecting alternatively spliced tandem repeats already known to be functionally and clinically important, and going beyond this current knowledge by identifying new repeats. Limitations of the present analysis include the focus on humans, the dependence on experimentally resolved PDB structures covering the repeats, and the fact that it did not account for interactions with nucleic acids. Future works will aim at overcoming these limitations. Another future direction will concern the development of integrated frameworks and learning algorithms to leverage these curated data for targeting protein interactions.

## Acknowledgments

This work benefited from computing resources funded by a grant of the French national research agency (MASSIV project, ANR-17-CE12-0009). The ANR-20-CE45-0010-01 RoDAPoG and the Hasso Plattner Institute provided a salary to A.S. Thanks to F. Oteri and H. Ripoche for their invaluable help with the computing infrastructure. Thanks to the Inria TAU team of the University of Paris-Saclay.

## Author contributions

Elodie Laine: Conceptualization, Methodology, Software, Validation, Formal analysis, Writing - Original draft preparation, Visualization, Supervision, Funding acquisition. Hugues Richard: Conceptualization, Methodology, Validation, Formal analysis, Data Curation, Writing - Review and editing, Visualization, Supervision, Funding acquisition. Antoine Szatkownik: Methodology, Software, Validation, Formal analysis, Investigation, Writing - Original draft preparation, Visualization. Diego Javier Zea: Software, Writing - Review and editing.

## Notes

### Competing Interest Statement

The authors have declared no competing interest.

### Summary of Updates

Panel B was aWe added a panel B to Figure 1 to better illustrate the method. We added new analyses regarding the evolutionary conservation of the repeats alternative usage (new subsection in the Results and Discussion). We emphasised more the biological findings of the study, with better explanations of the showcase examples. We rewrote the conclusion.

