## Supplementary File for "Building alternative splicing and evolution-aware sequence-structure maps for protein repeats"

### Supplemental Material

**Antoine Szatkownik[1,2+], Diego Javier Zea[3], Hugues Richard[1,2*], Elodie Laine[1*]**

[1] Sorbonne Université, CNRS, IBPS, Laboratoire de Biologie Computationnelle et Quantitative (LCQB), 75005 Paris, France. [2] Bioinformatics Unit, Genome Competence Center (MF1), Robert Koch Institute, 13353 Berlin, Germany. [3] Université Paris-Saclay, CEA, CNRS, Institute for Integrative Biology of the Cell (I2BC), 91198, Gif-sur-Yvette, France.

+ current affiliation: Université Paris-Saclay, CNRS, INRIA, LISN, Gif-sur-Yvette, France

### Supplementary Methods

#### Definitions

An ESG, introduced in (Zea et al., 2021), is a directed graph $\mathcal{S} = (\mathcal{V}, \mathcal{E})$, where each node $v \in \mathcal{V}$ is a spliced-exon (s-exon) and represents a multiple sequence alignment (MSA), $\tilde{s} = \{s_1, s_2, ...\}$, of translated exonic regions coming from a set of orthologous genes $G = \{G_1, G_2, \ldots, G_m\}$. Two nodes "start" and "stop" without sequences are added by convention. There is an edge in $\mathcal{E}$ from $v$ to $v'$ if there is at least one transcript coming from one species where the corresponding sequences in $v$ and $v'$ co-occur in a consecutive manner. A transcript corresponds to a path in the ESG from start to stop.

To study the alternative usage of s-exons, we define a canonical transcript, well-represented in all species, as detailed in (Zea et al., 2021). An AS event is then defined as a variation from this canonical transcript (**Fig. S1A**). More precisely, it corresponds to a pair of maximal subpaths that do not share any s-exon, where one subpath necessarily comes from the canonical transcript and the other one from some input transcript. We restrict the space of events by discarding variations between transcripts where none is a canonical transcript. Formally we define an AS event as a bubble in the ESG defined from the edge set $\mathcal{E}$. The event $b$ can be described as the tuple $((v_s, v_e), (v_1^c, v_2^c, ..., v_l^c), (v_1^a, v_2^a, ..., v_m^a))$,where each element is a list of nodes. The nodes $v_s$ and $v_e$ are the starting and ending anchors respectively. The list $(v_1^c, v_2^c, ..., v_l^c)$ defines the canonical subpath bounded by the anchors, while the list $(v_1^a, v_2^a, ..., v_m^a)$ defines the alternative subpath bounded by the anchors. Note that $l \geq 0$ and $m \geq 0$.

In the present work, in order to focus on the alternative usage of repeated protein regions, we introduce the notions of Alternatively Spliced Repetitive Unit (ASRU) and of spliced-repeat (s-repeat). An ASPR is a collection of s-repeats, where each s-repeat is an ordered set of s-exons. An ASPR must comprise at least two s-repeats. The s-repeats belonging to the same ASPR show substantial sequence similarity and are linked by at least one AS event, such that there is evidence of a conserved modulation of these s-repeats in the input transcripts (**Fig. S1B**). We propagate sequence similarity by transitivity such that we ask each s-repeat in an ASRU to share substantial sequence similarity with at least one other s-repeat in the same ASRU. The s-exons comprising a s-repeat are always co-expressed together such that the topology of the ESG does not break s-repeats.

#### Detailed algorithm

ASPRING starts from an ESG computed by ThorAxe (Zea et al., 2021) and representing the transcript variability of a set of orthologous genes $G = \{G_1, G_2, \ldots, G_m\}$. The four main steps of ASPRING algorithm are detailed in the following (**Fig. 1A**).

1. **All-to-all comparison of s-exons.** We convert all the s-exons' MSAs longer than 5 amino acids from FASTA format to A2M format using the *reformat* function from HH-suite3 (Steinegger et al., 2019). Then, we build $n$ probabilistic models, namely profile hidden Markov Models (HMM) (Eddy, 1998), with the *hhmake* function from HH-suite3. We globally align all profile HMMs with one another using *hhalign*, thus resulting in $n(n-1)/2$ alignments. In turn, we build a similarity graph reflecting the pairwise similarities between the profiles.

2. **Identification of similar s-exon pairs.** The identification of similar s-exon pairs consists in filtering the edges of the similarity graph. By default, we consider two s-exons to be similar if:

   - the p-value of the profile HMM-HMM alignment is lower than 0.01,

- the percentage of consensus sequence identity is higher than 45%,
- the coverage of the alignment is at least 80% for the query or the target,
- both s-exons are at least conserved in two species.

All these parameters are customizable by the users. Moreover, we require that the two s-exons in a pair are linked by at least an AS event (**Fig. S1B**). We record the set of AS events associated to each valid pair.

3. **Clustering of similar s-exons.** Next, we detect the connected components in the similarity graph, using only the edges that passed all previous filters. Each connected component is a prototype of an ASRU and each s-exon is a seed for a s-repeat.

4. **Refinement of the s-repeats.** The final phase refines the s-repeat definition to make the ASRU more homogeneous in length (in aa). That is, we seek to extend each seed with adjacent s-exons as much as possible while complying with the topology of the ESG. In practice, the algorithm loops over all valid s-exon pairs, and for each pair, it looks at the associated alignment to evaluate whether each s-exon could be extended through its N-terminal or C-terminal extremity (**Fig. S9A**). Given an extendable seed and an extension direction, the algorithm searches for possible extensions as s-exons adjacent to the seed in that direction in the ESG. Among these candidate extensions, only those that are always co-expressed with the seed are considered as valid. In other words, there is no AS event conserved in at least two species that breaks the junction between the seed and its extension. We verify this condition by computing a truth table. By accounting for the whole topology of the ESG, namely all events in the graph, we guarantee that there is only at most one valid extension for a given seed and direction. If a valid candidate extension is found and is shorter than 5 amino acids, then we concatenate it directly to the seed. If it is longer than 5 amino acids, we first verify that it shares sufficient similarity with the seed according to the criteria listed in step 2 before concatenating it to the seed. By iteratively applying the extension procedure, the algorithm progressively builds the s-repeats, within each ASRU (**Fig. S9D**).

### Analysis of the ASRU evolutionary and structural properties

For these analyses, we adopted a human-centred perspective by taking the human exonic sequences as references. We discarded the s-exons not present in human. This filtering reduced the number of s-repeats from 5 014 to 4 330.

##### Detection of sequence signatures

On each s-repeats pair, we used the information coming from the profile-profile alignment with HH-suite 3 to annotate sequence signatures. Consensuses (match states in the profile-HMM) from each s-repeat are first projected on the corresponding sequences in *H. sapiens* (note that this implies that sequences with no human representative are not conserved). According to the consensus, each position is either conserved or not conserved (annotated with a ~ character in the HHM-profile). Then, each position in the profile-profile pairwise alignment can be considered as similar (alignment score above 0.5) or not (score below $-0.5$). Based on those two pieces of information, we define three types of signatures. Specificity Determining Signature of type I (SDS type-I) occurs when only one of the two s-repeat positions is conserved. Specificity Determining Signature of type II (SDS type-II) occurs when both of the s-repeat positions are conserved, but the aligned positions are not similar. Finally, covariation corresponds to positions that variable in both s-repeats, but where the alignment is similar. To avoid biases due to the projection on one species, we additionally require that the proportion of the majority amino acid is below 0.75 in at least one of the pairs. In order to compare to a baseline of conservation, we also consider "Conserved match" positions. Those correspond to positions where both consensuses are conserved and the aligned consensuses are similar.

##### Disorder prediction

We predicted the content of intrinsic disorder in the s-repeats using the very rapid state-of-the-art method SETH (Ilzhoefer et al., 2022). It takes as input a single protein sequence and leverages embeddings computed from the protein Language Model ProtT5 (Elnaggar et al., 2021). It was trained in a supervised way against CheZOD scores, the latter being calculated from the difference between chemical shift values obtained by NMR spectroscopy (Howard, 1998) and computed random coil chemical shifts (Nielsen and Mulder, 2020). We fed SETH with all s-repeat containing transcript's sequences. Dealing with full-length transcripts allows accounting for the whole and possibly multiple sequence contexts of the s-repeats. We ran SETH on at total of 3 627 transcripts, among which 41 could not be treated because they were longer than 4 700 residues. As a result, we obtained disorder predictions from SETH for a total of 4 152 s-repeats. Following (Nielsen and Mulder, 2016), we considered a residue was disordered if its predicted CheZOD score was inferior to 8. We computed the proportion of disordered residues for each s-repeat averaged over the associated transcripts.

##### Structural coverage of interactions

We retrieved the list of PDB chains (PDB snapshot as of January 2023) associated with the ASRU-containing human genes from the web-based data mining tool BioMart (Smedley et al., 2009). We then applied the protocol described in (Dequeker et al., 2019) to identify the residues interacting with protein partners. More specifically, given a PDB chain, we retrieved all physiological protein-protein 3D complex structures available from the PDB where this chain, or a close homolog sharing > 80% sequence identity, was in contact with some other protein chain. We detected the interfacial residues with INTBuilder (Dequeker et al., 2017) and we mapped them back to the original chain through global pairwise sequence alignment (using the BLOSUM62 matrix, with the Biopython package (Cock et al., 2009)). A residue is considered at the interface if it is located at less than 5 Å from any atom from the partner. To determine the sets of homologs, we clustered the whole PDB using MMseqs2 (Steinegger and Söding, 2017) into groups of protein chains sharing more than 80% sequence identity and more than 80% sequence coverage.

We relied on the isoform.io resource providing 3D AlphaFold-predicted models for over 200 000 human transcripts (Sommer et al., 2022) to project the experimental structural information about interactions onto the s-exons. Each 3D model in the database corresponds to a transcript annotated in CHESS 3 (Varabyou et al., 2022). The database gives the mapping between the Chess IDs and the Transcript Ensembl IDs. We considered only the CHESS transcripts associated with an Ensembl transcript sharing the same sequence. We disregarded cases where the sequences did not have the same length or shared less than 95% of their residues in common. For each ASRU-containing gene, we superimposed the corresponding set of 3D models from isoform.io onto each of the PDB chains representing the gene using the *align* command from PyMOL API 2.5+ (DeLano, 2002). We then used the generated alignments to map the information about interacting residues onto the human sequences of the s-exons and s-repeats of interest. We labelled the residues resolved in the PDB but not in interaction with 0 (non-interacting), the ones resolved and found at interfaces with 1 (interacting), and the ones not resolved in the PDB with 2 (no information).

#### Supplemental Tables and Figures

Supplemental Table S1: **Identifiers of some genes and isoforms mentioned in this study.**

| Name | Ensembl gene ID | Ensembl transcript ID | Protein residue span | Figure |
|---|---|---|---|---|
| Matrilin 2 | ENSG00000132561 | ENST00000254898 | 285-647 | 1.B |
| Nebulin | ENSG00000183091 | | | |
| CPSF4 | ENSG00000160917 | | 1-116 | 3.C |
| ITGB1 | ENSG00000150093 | | 750-789 | 3.D |
| MAPT | ENSG00000186868 | | | |
| Ankyrin 1 | ENSG00000029534 | ENST00000347528 | 1-795 | 5.A |
| Tankyrase-1 | ENSMUSG00000031529 | ENSMUST00000033929 | 174-937 | 5.B |
| LGR5 | ENSG00000139292 | ENST00000266674 | 1-482 | 5.C |

The Ensembl gene and transcript identifiers are from version 105.

Supplemental Table S2: **Per-gene statistics computed over all human protein coding genes**

| **Variable** | **Mean** | **Std** | **Min** | **q25** | **Median** | **q75** | **Max** |
|---|---|---|---|---|---|---|---|
| Species | 8.9 | 2.7 | 1 | 8 | 10 | 11 | 12 |
| Transcripts | 19.7 | 12.7 | 1 | 11 | 17 | 26 | 132 |
| Events | 4.3 | 6.3 | 0 | 0 | 2 | 6 | 98 |

ThorAxe considers events supported by at least two transcripts.

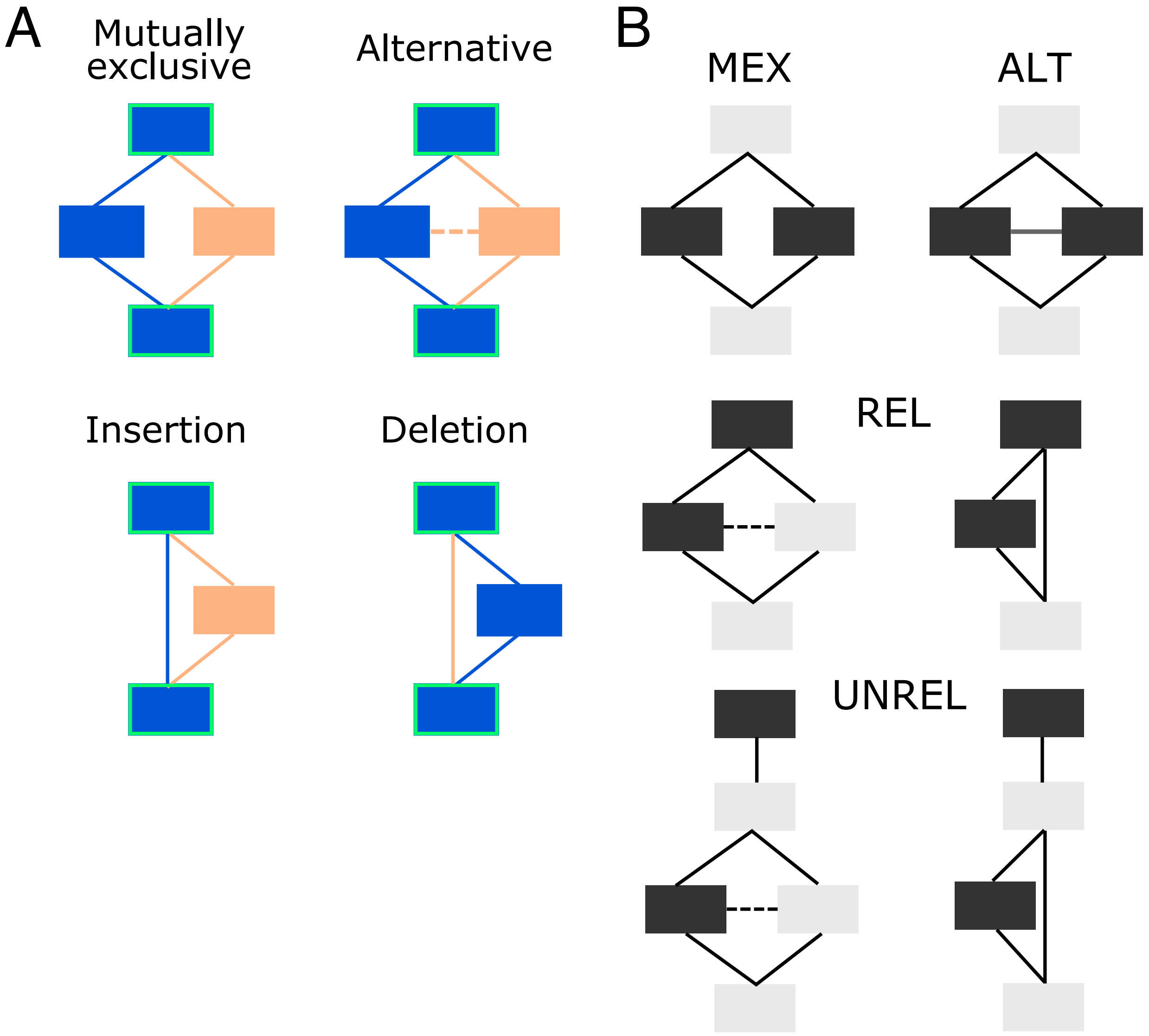


Supplemental Figure S1: **Alternative usage of s-exons.** **A.** Classification of AS events. The s-exons and edges present in the canonical transcript are colored in blue, while the alternative ones are in orange. An event is classified as mutually exclusive only if the two corresponding subpaths never co-occur in a transcript. Otherwise, it is considered as alternative. The green contours indicate that the s-exons anchoring the bubble may be absent, in the case of alternative initiations or terminations. **B.** Classification of similar s-exon pairs, highlighted in black. MEX: mutual exclusivity. ALT: alternative (non mutually exclusive) usage. REL: one s-exon is in the canonical or alternative subpath of an event (of any type), while the other one serves as a "canonical anchor" for the event. UNREL: one s-exon is in the canonical or alternative subpath of an event (of any type), while the other one is located outside the event in the canonical transcript. Each detected pair is assigned to only one category with the following priority rule: MEX>ALT>REL>UNREL.

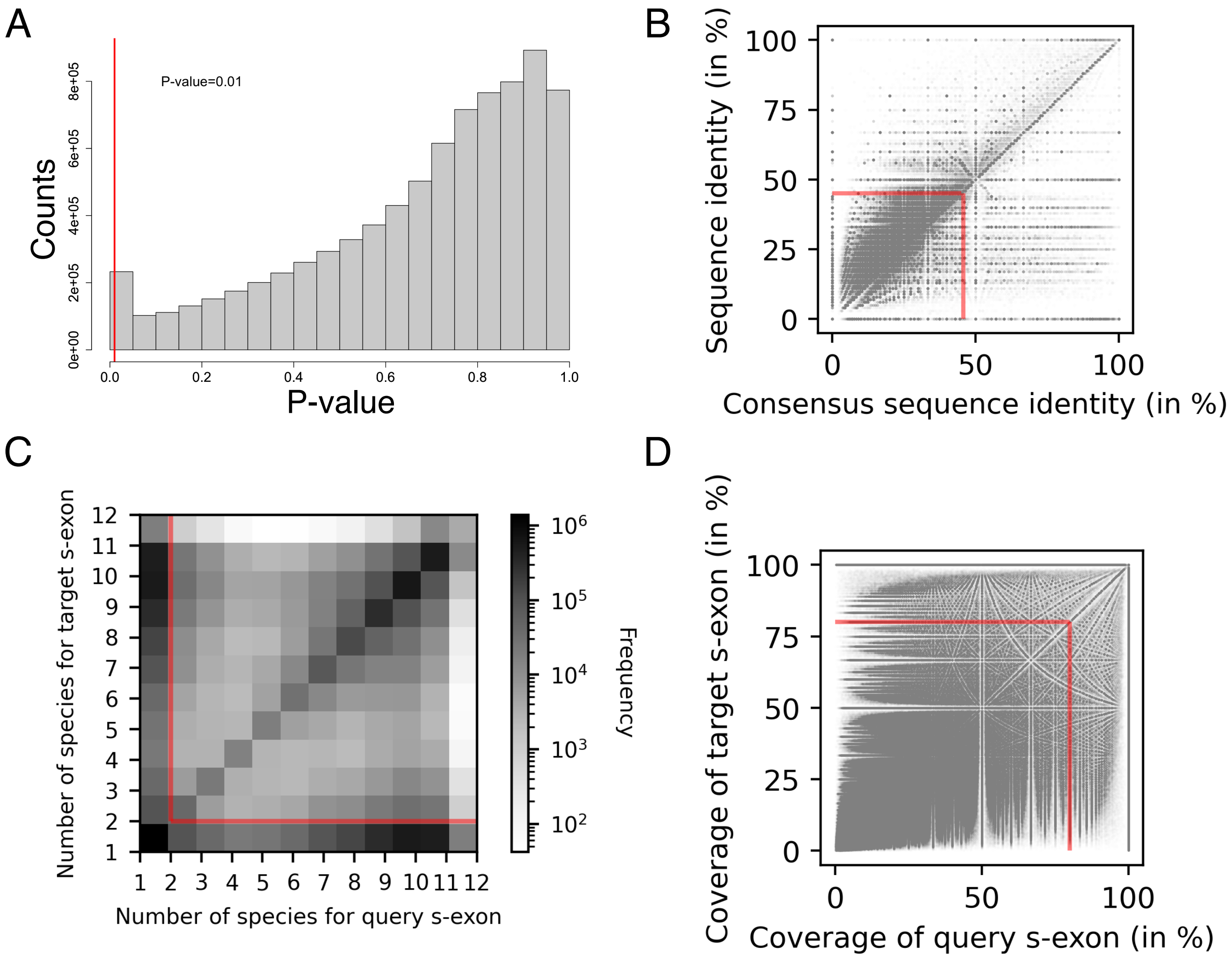


Supplemental Figure S2: **Influence of ASPRING filtering criteria.** We report the properties of an ensemble of 8 083 167 s-exon pairs extracted from our proteome-wide analysis across 12 species, from human to nematode. **A.** Distribution of the p-values, with only 2% of the pairs (158 775 pairs) displaying a significant p-value ($\leq 0.01$, on the left of the red line). **B.** Percentage of identity computed between the human sequences in function of the pHMM consensus sequence identity. Each dot represents an s-exon pair. About 14% of the pairs (1 126 772 pairs) are highly similar (>45% sequence identity, the dots outside the red square). **C.** Conservation levels of the two s-exons in each pair, measured as the number of species in the associated MSAs. In almost 40% of the pairs (3 037 220 pairs), the two s-exons are conserved in at least two species (delineated by the red square). **D.** Percentage of residues from each s-exon in a pair covered by the pHMM alignment. About 36% of the pairs (2 901 472 pairs) have their two s-exons covered at more than 80% (dots on the top and on the right of the red lines).

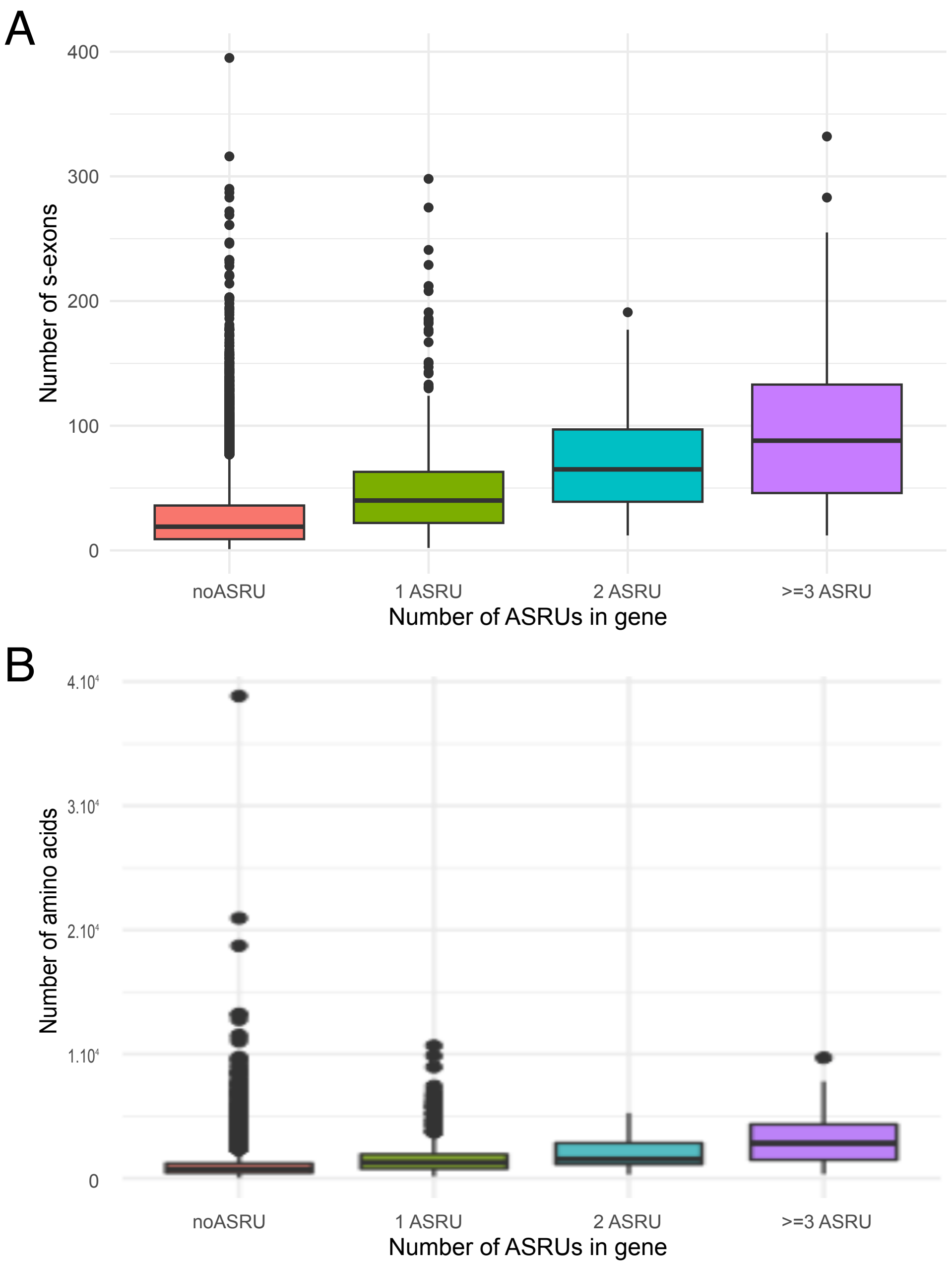


Supplemental Figure S3: **Distributions of gene length in function of the number of detected ASRUs.** The length is measured in s-exons (A) or in amino acids (B).

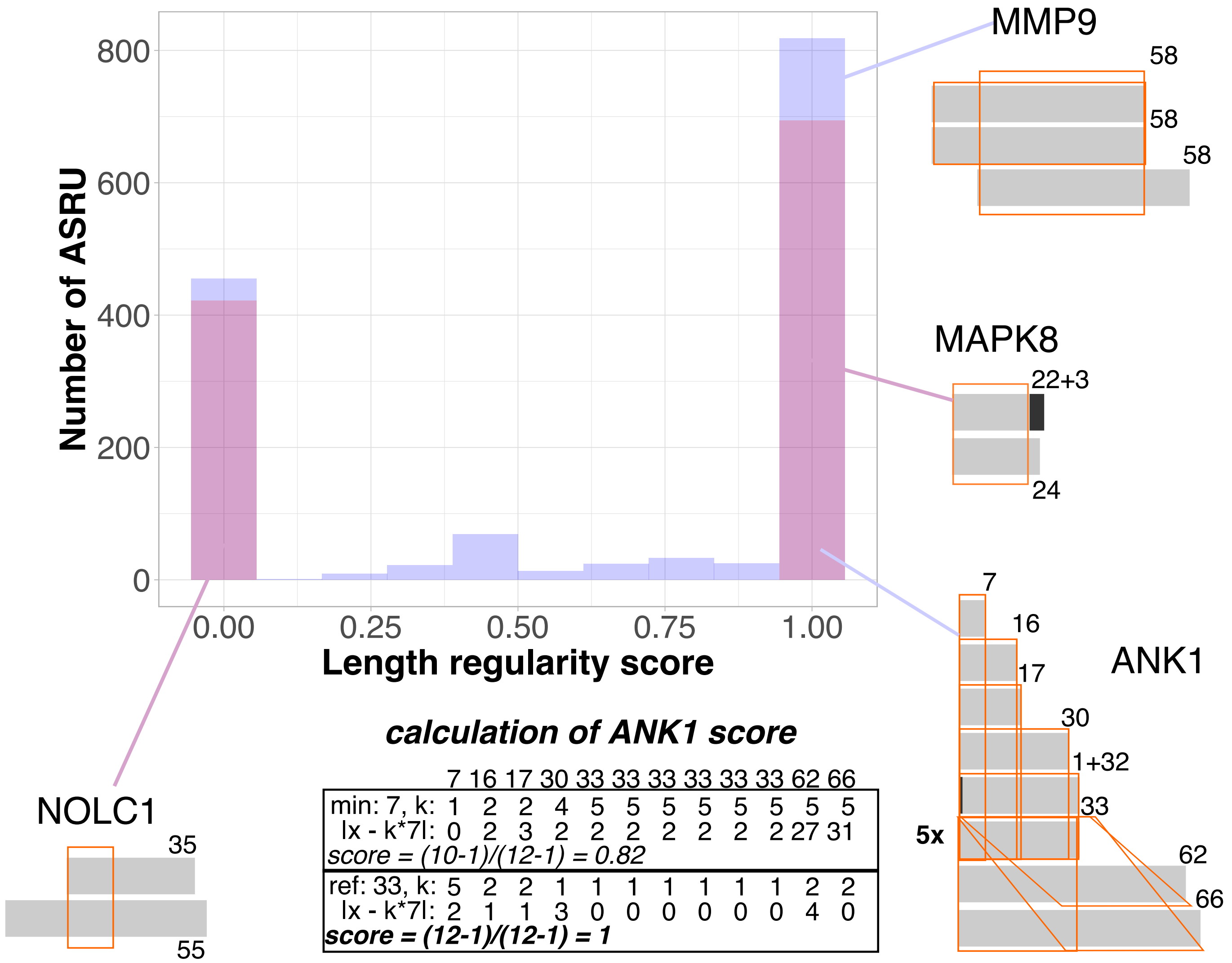


Supplemental Figure S4: **S-repeat length regularity.** The length regularity score reflects the extent to which the s-repeats have the same lengths or lengths that are multiples of each other within an ASRU. For each ASRU, we chose a reference s-repeat, of length $l_{ref}$, and we verified whether the other s-repeats were $k$ times longer or shorter, with $k = 1, 2, 3, 4$ or 5, with a tolerance of 5 amino acids. A score of 1 indicates that all s-repeats, excluding the reference, satisfy this condition. For the ASRUs comprising only two s-repeats (highlighted in pink), we took the length of the smallest s-repeat as reference. When dealing with the ASRUs containing more than 2 s-repeats, we considered the smallest length, and also the biggest length among the 50% smallest s-repeats. The schema around the plot show four illustrative examples, where each line corresponds to a s-repeat. When a s-repeat is made of several s-exons, the latter are colored in different grey tones. The lengths of the s-repeats (in amino acids) are indicated on their top right. The orange parallelepipeds delineate the aligned regions. We also give the detailed calculation of the score for the ASRU detected in ANK1. Please notice that for this ASRU, 5 s-repeats of the same length are represented by only one rectangle preceded by ”5x” for ease of visualisation.

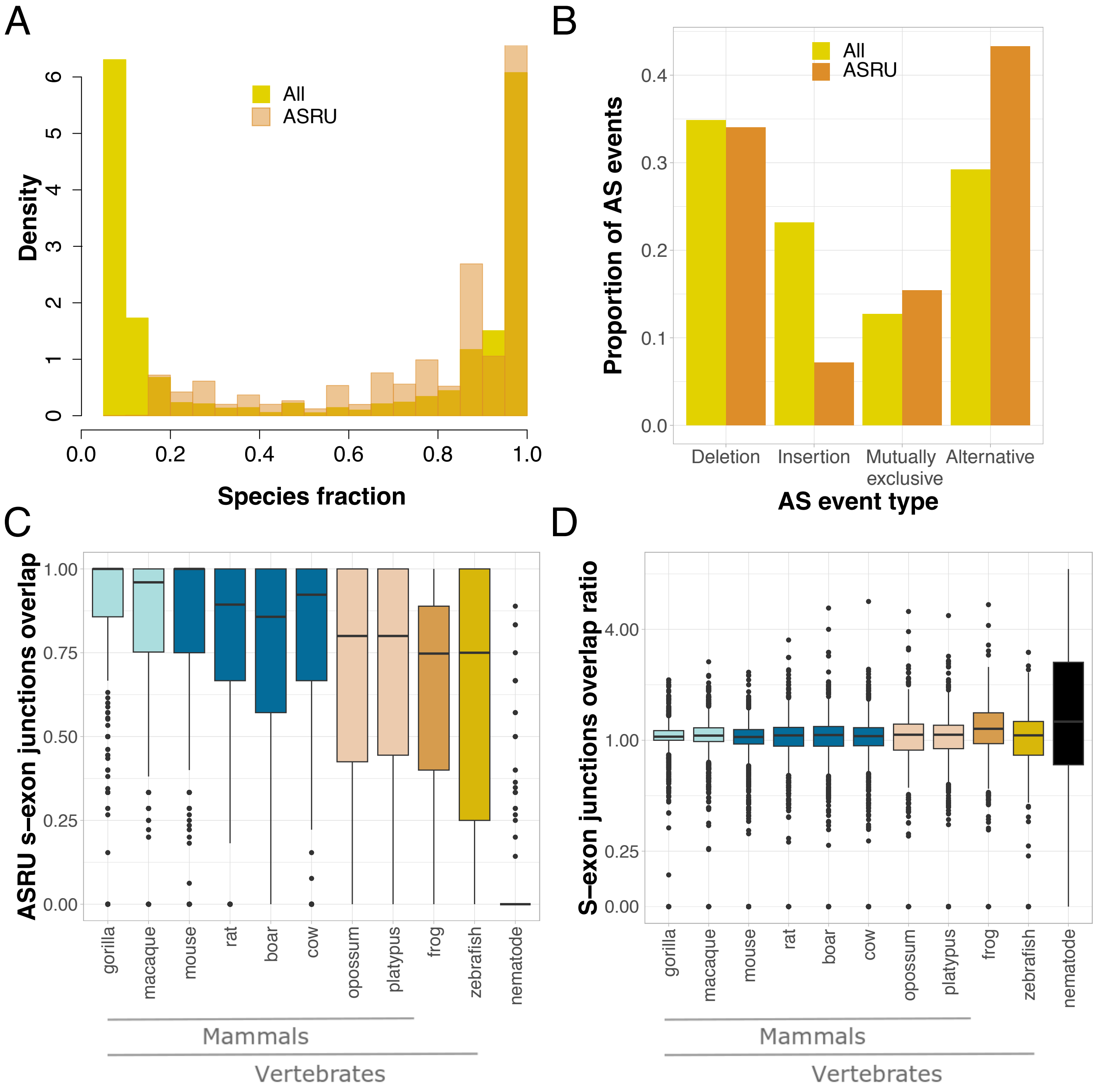


Supplemental Figure S5: **S-repeats evolutionary conservation and alternative usage. A-B.** We compare the distributions obtained for all s-exons over the whole human coding fraction (*ALL*, in yellow) with those corresponding to only the s-exons involved in the detected ASRUs (*ASRU*, in orange). **A.** S-exon conservation measured by the species fraction. The latter is computed as the number of genes/species where the s-exon is present, over the total number of orthologs considered. **B.** AS event types, as described in Fig. S1. **C-D.** S-exon junction overlap between human and each of the other species, focusing on the ASRU-containing genes. Each dot within each distribution corresponds to an ASRU. Given two sets of junctions $\mathcal{A}$ and $\mathcal{B}$, the overlap is computed as $\frac{2\times|\mathcal{A}\cap\mathcal{B}|}{|\mathcal{A}||\mathcal{B}|}$. **C.** Distributions of the overlaps computed for the junctions involving the detected s-repeats. **D.** Distributions of the overlap ratios between the s-repeat junctions and all junctions in the ESG.

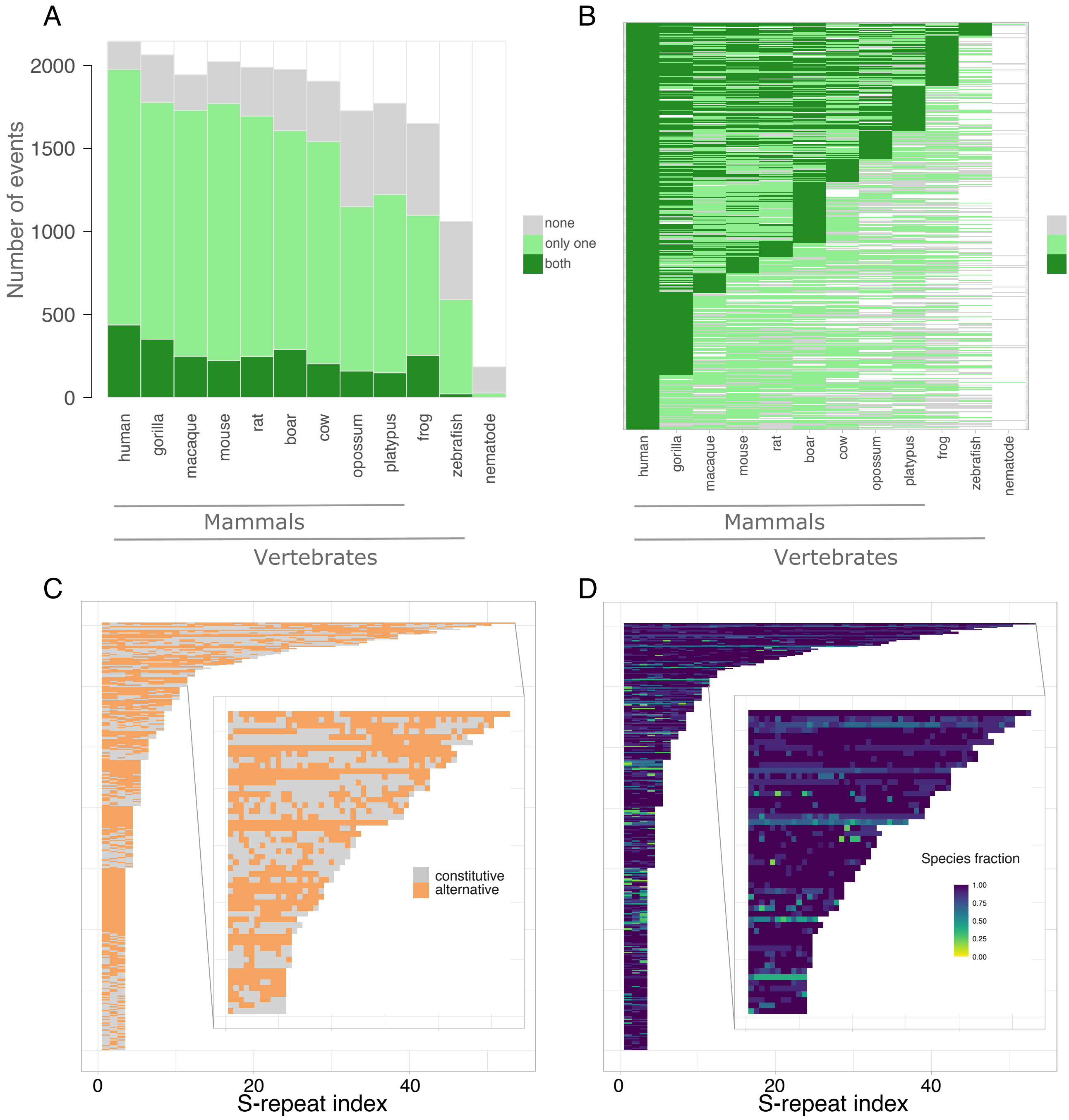


Supplemental Figure S6: **Landscape of the alternative usage of the detected s-repeats in evolution. A.** Supporting evidence for the 2 555 events associated with the 1 469 detected ASRUs. In the ESG, an event corresponds to a pair of subpaths, one being canonical and the other alternative. Within each species, we report the number of events for which both subpaths are present (dark green), only one subpath (light green) or none (grey). The white part indicates the amount of events that cannot be assessed because the corresponding human query genes do not have a one-to-one ortholog in the considered species. **B.** Conservation of the 489 events for which both the canonical and alternative subpaths are present in human. The color code is the same as in panel A. **C-D.** Heatmaps showing the alternative usage (C) and species fraction (D) of each s-repeat within each of the ASRUs containing more than 2 s-repeats. The insets zoom on the ASRUs with more than 10 s-repeats. A constitutive s-repeat is part of the canonical transcript and always included in the transcripts. An alternative s-repeat takes part in the canonical of the alternative subpath of at least one event. We omitted the longest ASRU, coming from Nebulin and containing 127 s-repeats, among which 115 are alternative, for ease of visualisation.

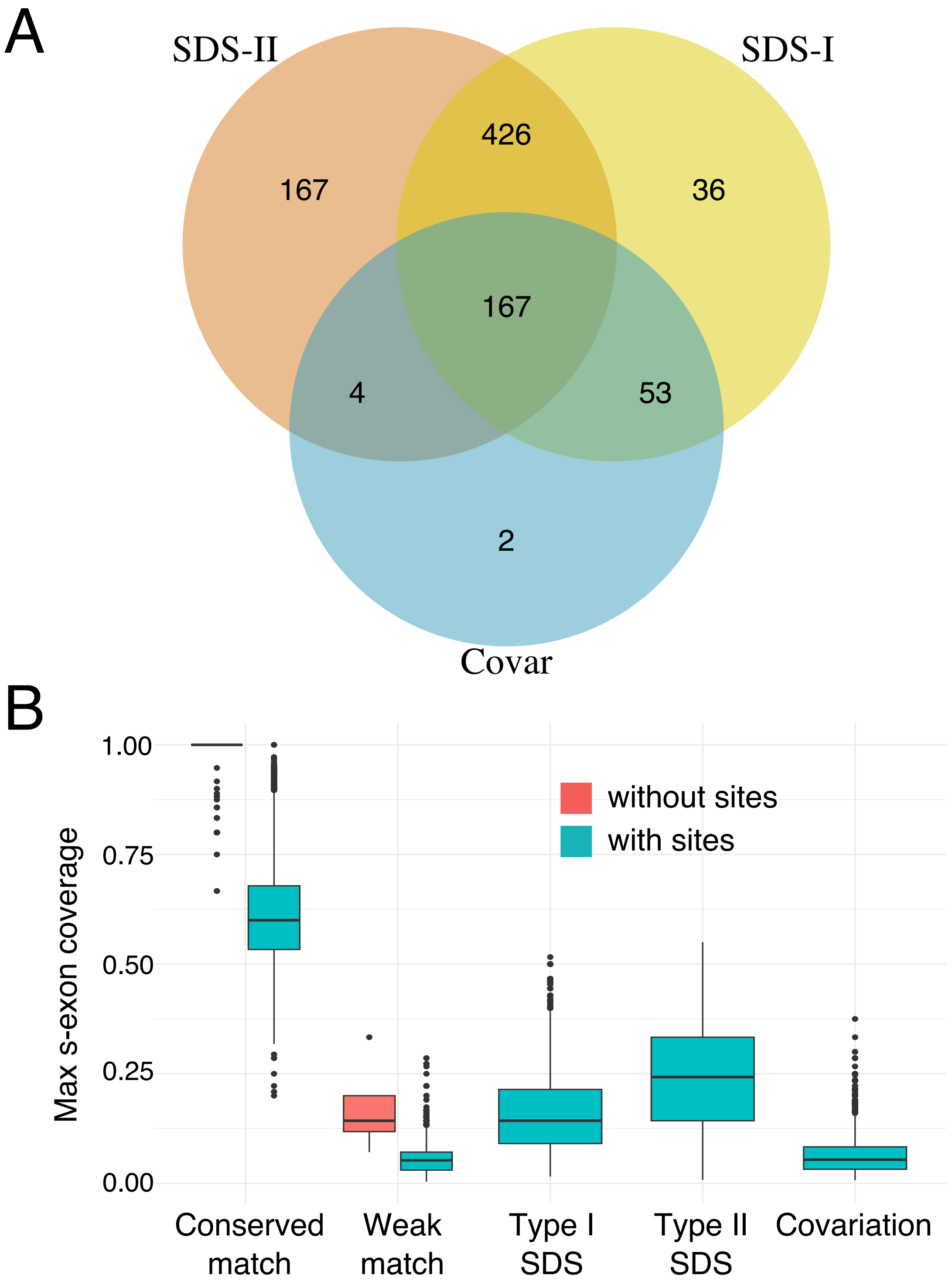


Supplemental Figure S7: **Specificity determining and covariation sites.** **A.** Venn diagram reporting the number of ASRUs where at least one pair of s-repeats has specificity determining sites (SDS) of type I or II, or covariation sites. **B.** Comparison of the distributions of the proportion of aligned positions displaying different characteristics in the s-repeats pairs with SDS and covariation sites (in turquoise) versus those without (in salmon). A conserved match is a perfect match in the HH-suite nomenclature (column score above 1.5, conserved consensus) and a weak match is a semi-positive match (column score between -0.5 and 0.5). The proportion is computed as the fraction of the number of positions complying with the criterion of interest over the length of the smallest s-exon in the pair.

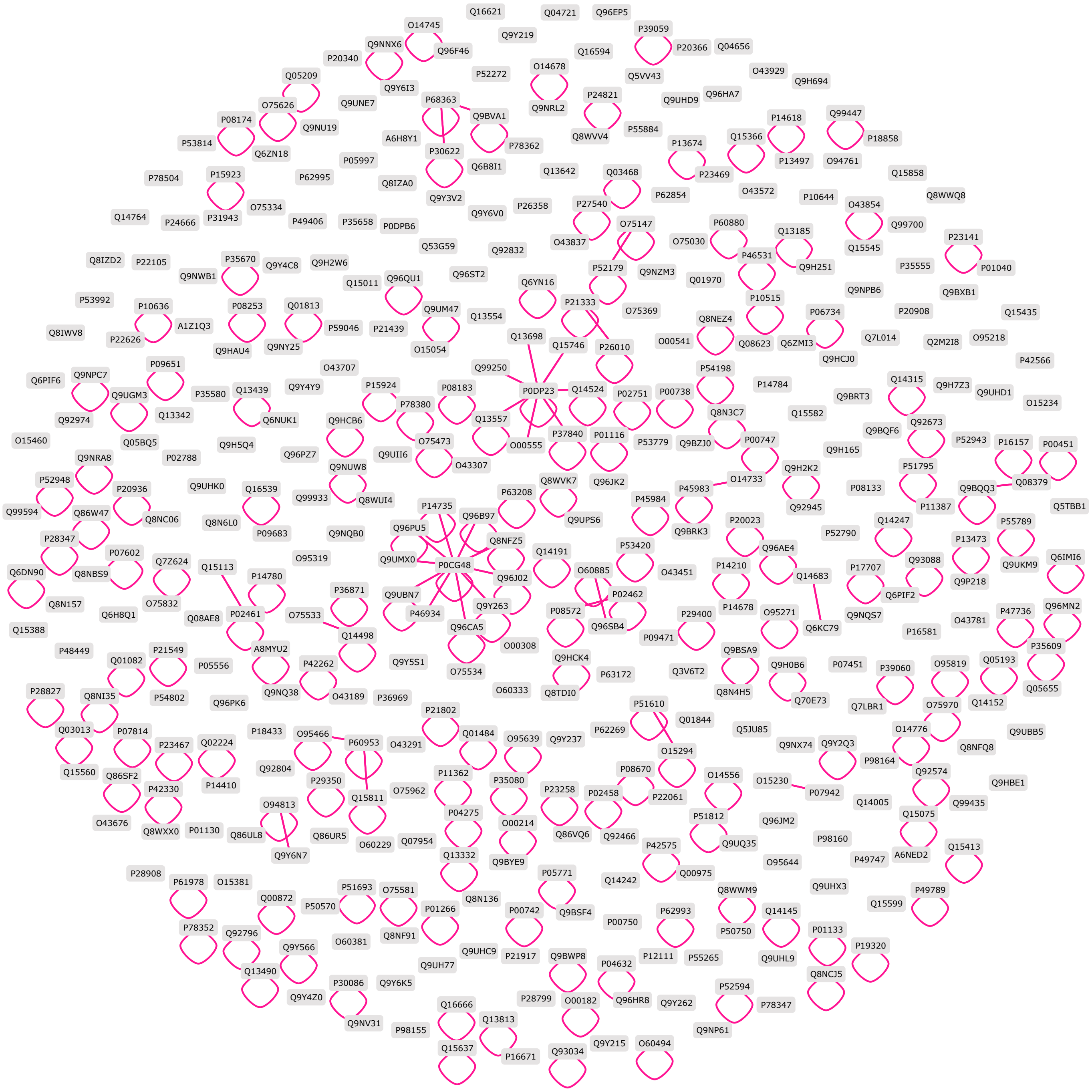


Supplemental Figure S8: **Structurally resolved 3D interaction network among the ASRU-containing proteins.** Each node represents an ASRU-containing gene and is labelled with its Uniprot ID. An edge between two nodes indicates that the corresponding proteins or their very close homologs at 95% sequence identity interact with one another in a physiologically relevant 3D protein complex structure available from the PDB. The supporting 3D complex structures are likely to involve only one main isoform for each protein, since the whole PDB contains very few alternative isoforms (Ait-hamlat et al., 2020). The much higher number of homo-dimers compared to hetero-dimers reflect a general tendency in the PDB (Bertoni et al., 2017). The network was inferred and rendered with LEVELNET (Behbahani et al., 2023).

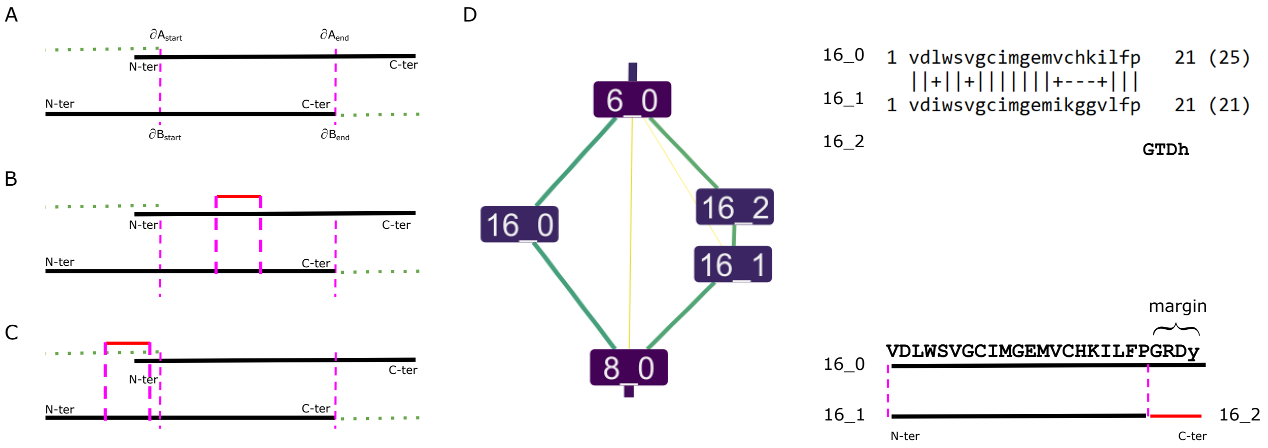


Supplemental Figure S9: **Schematic illustrations of the extension algorithm. A.** Aligned pair of s-exons $A$ and $B$. The boundaries of the alignment are indicated by dashed pink lines. The s-exon $A$ could be extended through its N-terminus while the s-exons $B$ could be extended through its C-terminus (see the green dotted lines indicating the margins). **B.** The alignment of the candidate extension $C$ (in red) with the s-exon $B$ overlaps with the alignment between the s-exons $A$ and $B$. In that case, the extension is rejected. **C.** Case where there is no overlap and the candidate extension is retained.**D.** Example of an ESG where the s-exons $16_0$ and 16_1 form a valid similar s-exon pair. While all the amino acids from 16_1 are present in the alignment, only 21 out of 25 amino acids from 16_0 have been aligned. Hence, the s-exon 16_1 could be extended through its C-terminus. The candidate extension 16_2 is valid according to the ESG topology and it is shorter than 5 amino acids, so it can be concatenated to 16_1. This ESG will thus lead to the identification of one ASRU comprised of two s-repeats, one made of a single s-exon (16_0) and the other made of two s-exons (16_1 and 16_2).
